# Restoring signatures of consciousness by thalamic stimulation in a whole-brain model of an anesthetized nonhuman primate

**DOI:** 10.1101/2025.11.07.686535

**Authors:** Eider Pérez-Ordoyo, Jordy Tasserie, Lynn Uhrig, Chloé Gomez, Antoine Grigis, Wiep Stikvoort, Sebastian M. Geli, Enzo Tagliazucchi, Morten L. Kringelbach, Andrea I. Luppi, Gustavo Deco, Béchir Jarraya, Yonatan Sanz Perl

## Abstract

Treatment options for Disorders of Consciousness (DoC) are limited due to insufficient understanding of the underlying neurobiological mechanisms. Two primary strategies for characterizing DoC and assessing treatment efficacy are *in vivo* experiments with animal models, and *in silico* computational models. We combined both approaches by creating a whole-brain model tailored to the experimental functional magnetic resonance imaging (fMRI) data of a single anesthetized macaque. It was previously reported in an *in vivo* experiment that anesthesia-induced loss of consciousness was partially reversed by specific electrical stimulation of the thalamic central nuclei. The *in silico* model reproduced the brain dynamics underlying the restoration of consciousness, providing a potential explanation for the transition between these brain states as continuous trajectories unfolding in a low-dimensional space. Our results demonstrate that whole-brain computational models reproduce the spatiotemporal properties of fMRI recordings during loss of consciousness and during its recovery induced by electrical stimulation, enabling computational exploration of perturbation-based interventions to potentially personalize treatment and aid recovery of consciousness in DoC patients.

## Introduction

Loss of consciousness occurs as a consequence of neurobiological processes during sleep, pharmacological interventions in anesthetized individuals, and pathological processes in brain injured patients diagnosed with Disorders of Consciousness (DoC). A fundamental problem in the study of consciousness is the need to characterize brain states that differ in their first-person experience using objective data and metrics. Different theoretical developments, such as the theory of recurrent thalamo-cortical loops [1], the mesocircuit hypothesis [2], the Integrated Information Theory (IIT) [3], dendritic integration theory [4], and the Global Neuronal Workspace (GWT) [5], have been proposed to inform the relationship between consciousness, brain activity, and its capacity for information processing. In spite of intensive investigation, the brain mechanisms underlying global states of consciousness remain elusive, hindering the adequate diagnosis of brain-injured patients with DoC and severely limiting the available options for their treatment.

The use of neuroimaging and electrophysiological data has enabled the development of data- and theory-driven signatures of consciousness beyond behavioral clinical scores such as the Glasgow Coma Scale (GCS) [6], [7] and the Coma Recovery Scale-Revised [7], [8], [9] (CRS-R). While these behavioral scales are frequently used for the diagnosis and prognosis of DoC patients, a key limitation is their reliance on overt motor or verbal responses. However, unresponsiveness is not synonymous with unconsciousness. Under anesthesia and in sleep, conscious experience can persist despite unresponsiveness (in dreams for example), highlighting the conceptual differences between the two [10]. Evidence from neuroimaging further shows that some behaviourally unresponsive patients retain volitional cognition: previous fMRI work demonstrated the ability to follow commands via mental imagery in a patient meeting criteria for the vegetative state [11], and more recent data from different centers established the prevalence and prognostic relevance of cognitive-motor dissociation, a condition by which cognition is preserved with impaired motor output [12]. Furthermore, mechanistic reviews of DoC also emphasize that abnormalities in arousal, integration, or motor efference can mask awareness, motivating the use of neuroimaging techniques and perturbational approaches to assess capacity for conscious processing beyond the limits already mentioned of behavior [13]. In this context, perturbational approaches have the ability to probe the brain’s capacity to quickly adapt and re-organize when facing rapidly changing conditions, representing a signature of conscious states [14], [15]. Importantly, perturbative approaches can also be pursued to destabilize the brain activity patterns characteristic of DoC, with the objective of transitioning toward a state of conscious wakefulness [16], [17], [18]. This possibility is supported by computational modeling studies indicating that brain states are not only characterized by their compatibility with conscious content, but also by their stability against external perturbations [14], [18], [19], [20]. For instance, slow-wave sleep, as a state of reduced consciousness, can transition toward conscious wakefulness in response to weak external or endogenous perturbations [21], [22]. In contrast, inducing transitions toward consciousness in DoC patients is less straightforward and may require targeted interventions informed by disease mechanisms [23], [24], [25].

Previous clinical studies demonstrated that external brain stimulation can modulate the behavioral responsiveness in states of reduced consciousness, including DoC patients. Invasive electrical stimulation, such as the deep brain stimulation (DBS) technique, has provided encouraging results improving behavioral signatures of consciousness in DoC patients [26], [27]. Non-invasive electrical stimulation, such as transcranial direct current stimulation (tDCS), was also investigated as a potential avenue to accelerate the recovery of consciousness in brain-injured patients [23], [28], [29], [30]. Some of the most promising results arise from brain stimulation experiments conducted in anesthetized non-human primates as an animal model of unconsciousness. It has been shown that selective stimulation of the macaque intralaminar nuclei of the thalamus can restore signatures of consciousness in monkeys under anesthesia [31], [32], [33], [34], suggesting a potential causal role of the central thalamus in supporting consciousness. Moreover, brain stimulation also improved neuroimaging-based signatures of consciousness, such as decreasing the similarity between whole-brain functional connectivity (FC) and structural connectivity (SC).

Despite these promising results, *in vivo* experiments offer a limited perspective on how and where brain stimulation is capable of restoring consciousness; in particular, the systematic and exhaustive assessment of targeted perturbations is beyond the reach of animal studies. In this context, *in silico* ones are a cost- and time-effective alternative to explore how different forms of brain stimulation can induce transitions between healthy and pathological brain states [22], [35], [36]. The main goal of this work is to determine whether computational models of whole-brain activity can inform on the recovery of consciousness due to targeted stimulation. We leveraged a published dataset [34] where macaques under propofol anesthesia were stimulated using a DBS device targeting the central thalamic (CT) nuclei (CT-DBS) or a control structure, the ventrolateral (VL) thalamic nuclei (VL-DBS), providing evidence of the specificity and the efficacy of central thalamic DBS to restore consciousness. Here, we developed whole-brain computational models [37] fitted to individual macaques under different experimental conditions. Using these models, we generated low-dimensional representations of brain states through a Deep Neural Network architecture known as Variational Autoencoder (VAE) [38]. These low-dimensional representations enabled us to explore *in silico* perturbations to test whether we could match the simulated brain activity with that of the experimental stimulations, and to identify candidate targets capable of eliciting similar brain activity. Based on these findings, we proposed novel intervention targets based on literature defining the Global Neuronal Workspace network for macaques (GNW) [39]. We examine the theoretical predictions perturbing the GNW, showing that the cortical dynamics differ from those of the experimentally explored conditions.

## Results

### Methodological overview

We investigated how perturbations applied to a model of whole-brain activity could drive transitions from brain states representative of anesthesia toward conscious wakefulness. **Figure 1** contains an overview of the followed methodology, which is similar to that of past literature [16], [17], [38], [40]. The whole-brain model was informed by empirical data from an experiment investigating the effect of DBS to the Central Thalamic nuclei (CT-DBS) on behavioral and neural signatures of consciousness in an anesthetized non-human primate. The main result of this study was that CT-DBS delivered at 5V led to the recovery of signatures of consciousness.

**Figure 1.**
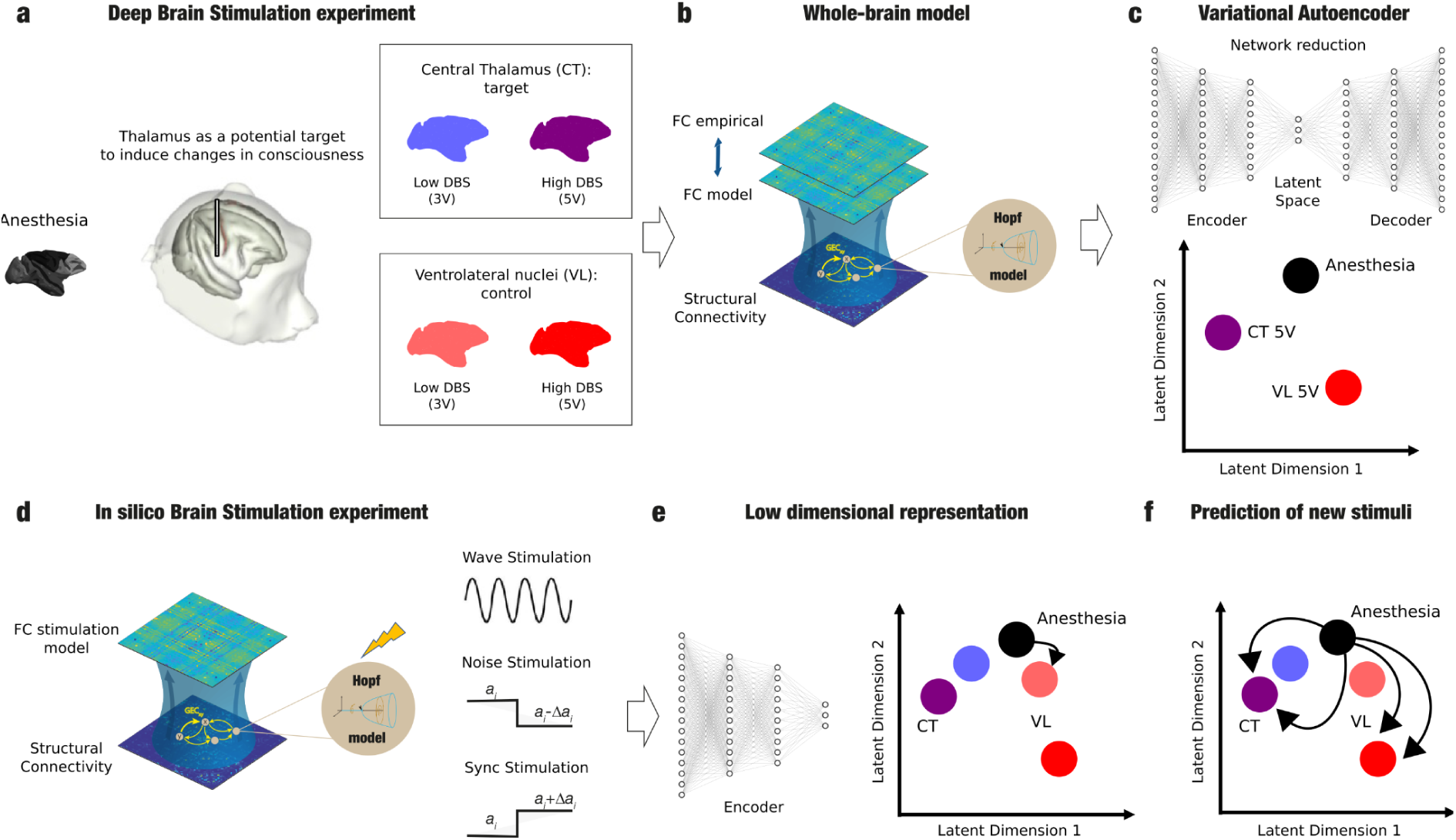
Methodological overview. Starting from the empirical data, we created a whole-brain model based on Hopf (or Stuart-Landau) nonlinear oscillators. The model was fitted to the empirical functional connectivity (FC). The simulated dynamics were then used to train a Variational Autoencoder (VAE), resulting in a 2-dimensional “latent space”, providing a straightforward geometric characterization of the different brain states. Finally, transitions between these brain states could be observed in the latent space through the application of different perturbations to the model.

Blood Oxygen Level-Dependent (BOLD) signals obtained from fMRI scans were parcellated according to the CoCoMac parcellation [41], comprised of 41 cortical regions per hemisphere, and used to compute whole-brain FC matrices for the following conditions anesthesia, CT stimulation (3 volts and 5 volts), and VL stimulation (3 volts and 5 volts). We used data from a single monkey to study transitions at the single-subject level, consistent with the need for personalized models of DoC patients (**Figure 1a**). Later, we used data from a second monkey to reproduce the findings from the first. Subsequently, we fitted a Hopf whole-brain model to the FC matrix obtained from each condition. This model consisted of nonlinear noisy Stuart-Landau oscillators which simulate the local dynamics of each brain region. These oscillators were coupled according to the brain’s SC, allowing the model to capture interaction between brain regions. In this model, the dynamics of an isolated node are governed by the normal form of a Hopf bifurcation, with the bifurcation parameter (*a*) defining qualitatively different regimes: self-sustained oscillations around a limit cycle for (*a > 0*), and a stable fixed point with noise-driven fluctuations for (*a < 0*)

To study how well the dynamics simulated by our model matched the empirical data, we quantified the goodness of fit (GoF) of the models by comparing the empirical and simulated FC (sFC) matrices, optimizing the model parameters to obtain the highest value of SSIM for each condition (**Figure 1b**). We quantify the similarity between empirical and simulated FC matrices using the Structural Similarity Index (SSIM), which is a known measurement for images taking into account luminance, contrast and structure. It has been previously used in the field to compare FC matrices because it offers a balance between absolute and relative differences and for its intuitive interpretation as a metric that factors in correlation and Frobenius distances [22], [38], [40], [42].

Our models were optimized using two steps, resulting in heterogeneous models as presented in Ipiña et al. [43]. To do so, we first optimized the models choosing the global coupling value that maximized the GoF. Afterward, we optimized the local dynamics of each pair of homotopic brain regions (tweaking the bifurcation parameter) to match the empirical static FC topography. Crucially, our heterogeneous models outperformed the homogeneous models (no bifurcation parameter tuning) across conditions, achieving an improved the GoF for all conditions, achieving a high fit to both spatial and dynamical properties of the living brain compared to previous work [36], [42], [43].

As a final step, we used simulations from our most accurate heterogeneous models as a data augmentation tool to train a Variational Autoencoder (VAE), a deep learning approach for nonlinear dimensionality reduction. This method enabled us to obtain a low-dimensional representation of the dynamics and to cluster states based on the dynamics’ underlying characteristics [36], [38]. Because deep learning requires substantially more data than what is available from a single subject, data augmentation allowed us to generate sufficient surrogate data that preserved the characteristic features of empirical dynamics while introducing sufficient variability to prevent overfitting [40]. The VAE learned a two-dimensional latent space that captured the structure of the surrogate FC matrices. This unsupervised representation revealed distinct clusters corresponding to experimental conditions, indicating that underlying features of brain dynamics alone were sufficient to separate states (**Figure 1c**) [36]. Assisted by this representation, we explored different plausible emulations of the stimulation protocol used in the experiment using different local perturbations, as done previously in the literature: wave perturbation (a periodic perturbation delivered at the dominant frequency of the local BOLD fluctuations), synchronizing perturbation (shift to a*>0)*, and desynchronizing or noise perturbation (shift to *a<0)* (**Figure 1d**) [16], [21], [22], [36]. We investigated whether our modeling approach could match the experimental results by encoding the FCs of the stimulated dynamics in a low-dimensional space and comparing them with the dynamics obtained by perturbing the anesthesia model targeting the brain regions activated in the *in vivo* experiment (**Figure 1e**). Finally, we exhaustively explored the repertoire of possible stimulations to determine new targets to displace the system’s dynamics as in the different conditions of the experiment (**Figure 1f**).

### Modeling the brain dynamics corresponding to different levels of consciousness

We followed a two-step model fitting procedure in order to fit single-subject whole-brain models for each condition. In the first step, we used a homogenous model (meaning that the bifurcation parameters of all the nodes in our model were fixed to the same value, in this case *a = -0.02*, as it had been proved to be the best at simulating resting state activity [42], [44]) and optimized the SSIM as a function of the Global Coupling parameter (G) (**Figure 2a**). We observed that the anesthesia condition had the optimal fit for a lower G value compared to the other conditions, being 5V CT-DBS the condition for which G was the highest, indicating that 5V CT-DBS induces greater synchronization and interconnectivity than anesthesia and the other conditions.

**Figure 2.**
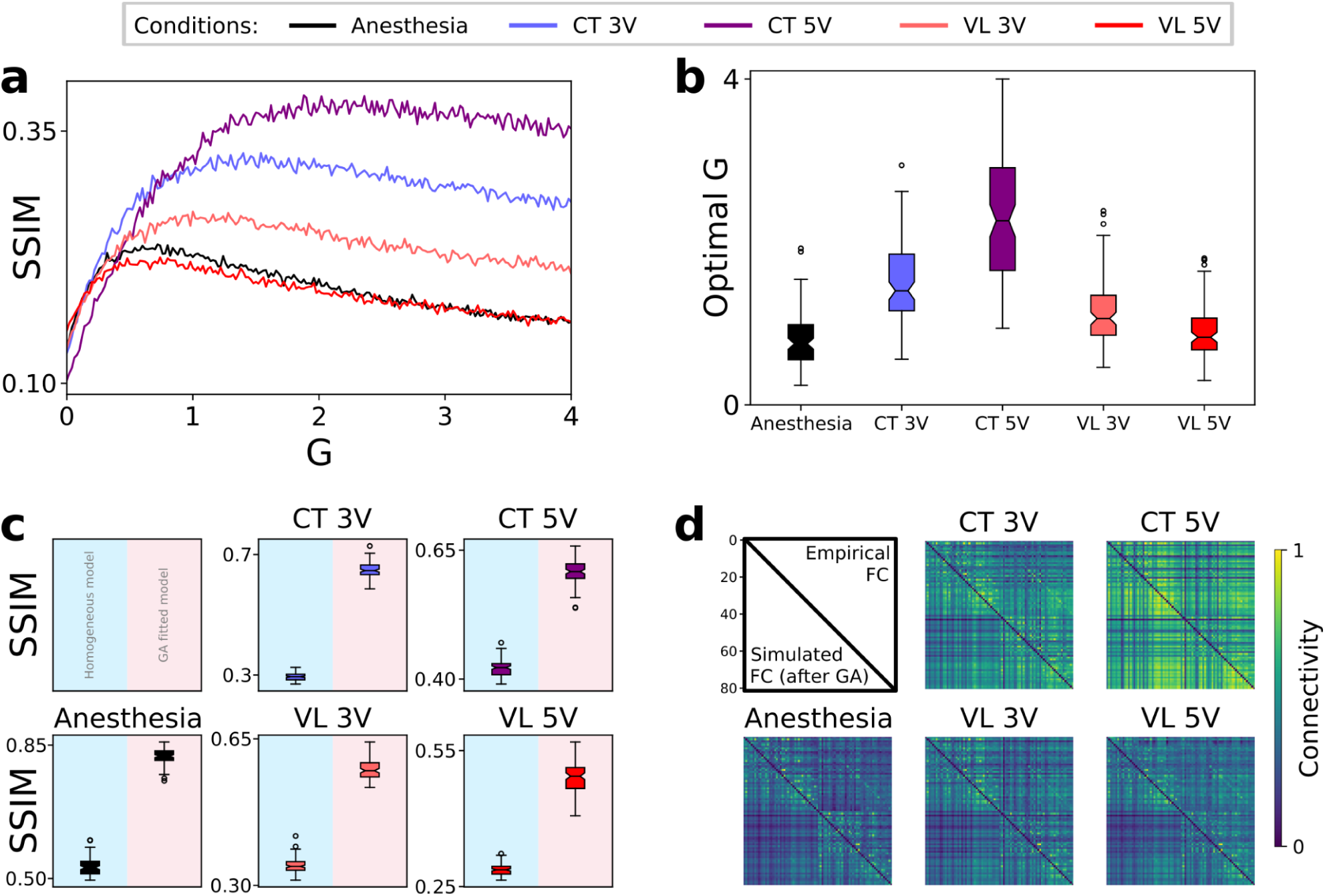
The whole-brain model reproduced experimental conditions for different choices of coupling and bifurcation parameters. (a) Curves of mean SSIM for homogeneous Hopf models (*a =-0.02*) as a function of the Global coupling parameters (G) for each experimental condition. (b) Optimal G distributions for different conditions. Each distribution was generated after a single exploration of the G parameter space. (c) SSIM values by condition, comparing the homogeneous vs. heterogeneous models. All the changes in SSIM between both versions of the models were significant (*p* < *0.001*). (d) Graphical representation of functional connectivity (FC) matrices, showing the empirical one on the upper triangle of the matrix and the simulated one on the lower triangle.

Next, we fitted the local bifurcation parameters. To simplify the procedure, we set identical parameter values for homotopic brain areas, assuming similar dynamics given similar cytoarchitectures across corresponding regions of both hemispheres. We fitted the resulting 41 bifurcation parameters using a genetic algorithm (GA) to find an optimal model for each condition. By performing this second step we generated a more complex model capable of capturing the variability of the dynamics across the cortex, which improved the goodness-of-fit significantly. The median of the SSIM values of our models in the GA improved by an average of 0.3 ± 0.1 in SSIM across all conditions. Some conditions improved more than others: the anesthesia condition’s SSIM median increased by an average of 0.42 ± 0.07 showing the highest improvement, while 5V VL-DBS showed the least improvement, increasing by only 0.14 ± 0.04 (**Figure 2c**).

### Low-dimensional representation of whole-brain dynamics

To investigate whether the distinct brain states elicited by thalamic DBS could be distinguished in a low-dimensional space, we trained a VAE using surrogate data from our most accurate heterogeneous models of the anesthesia, 5V CT-DBS and 5V VL-DBS conditions. These conditions were selected to evaluate whether the VAE could generalize to unseen data, in this case the lower-voltage conditions, under the hypothesis that these would be encoded between the anesthesia and the corresponding high-voltage states. These three training conditions were chosen to represent the full range of observed brain states. The resulting 2D latent space (**Figure 3a**) clearly separated the training conditions, and when encoding the remaining conditions with unseen surrogate data from the training conditions, the newly seen conditions occupied intermediate positions between the anesthesia state and their respective 5V counterparts (**Figure 3b**), consistent with our hypothesis. Using the decoder network, we decoded a grid of coordinate pairs *(z_1_, z_2_)* that covered an area of the latent space that extended from that of where the known dynamics lie. With this, we obtained a grid of FC matrices that correspond to different regions of the latent space. This allowed us to examine unexplored regions of the latent space and characterize the dynamics of the pathways through which perturbations induce state transitions in the brain, even when these transitional states were not captured in the experiment (**Figure 3c**).

**Figure 3.**
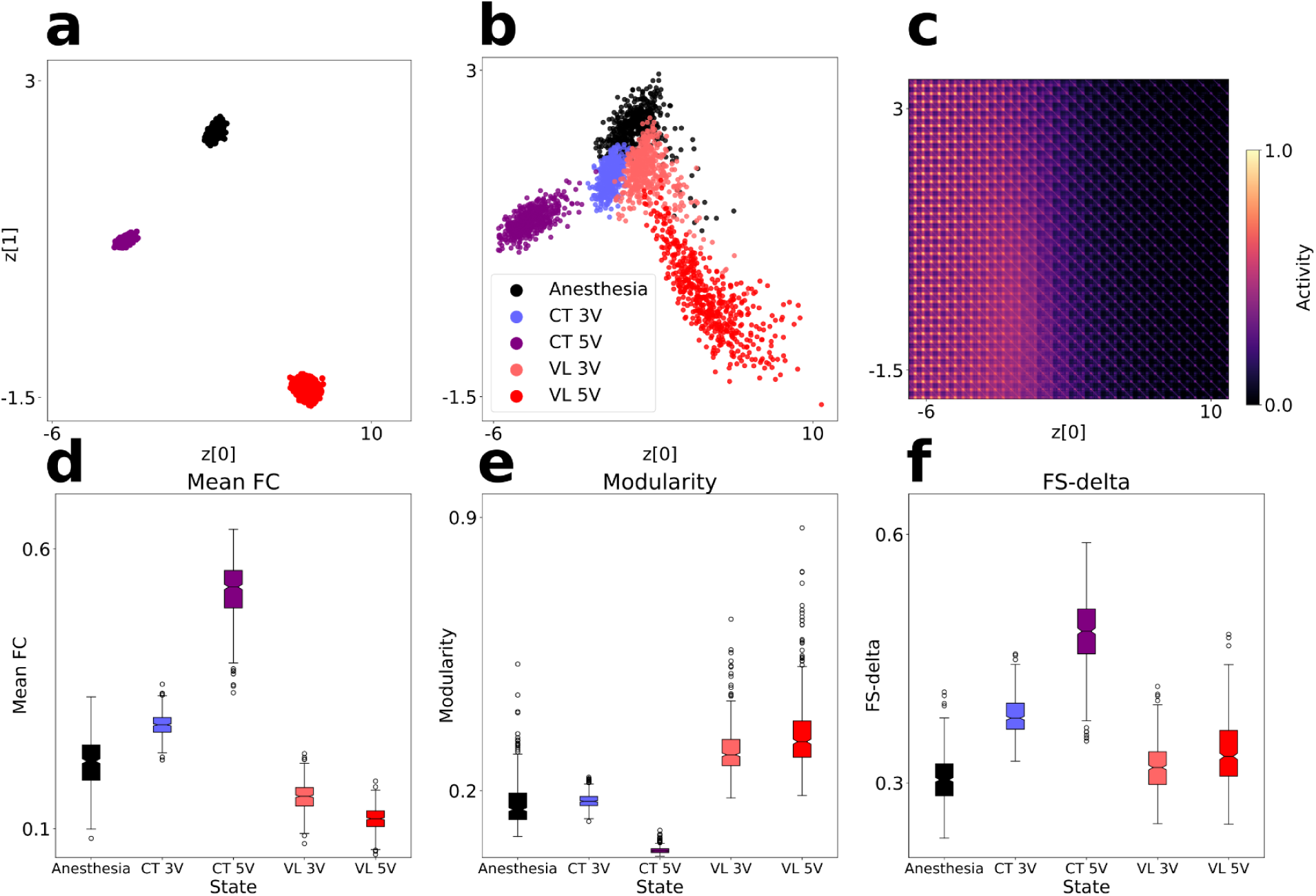
Latent space dimensions encode stimulation sites and signatures of consciousness. (a) Training points encoded into the latent space, with z[0] and z[1] being its latent space dimensions. (b) FCs previously unseen by the VAE (corresponding to all experimental conditions) encoded into the latent space. (c) Decoded FC matrices from a grid of points covering the latent space. Each FC is centered around the coordinates of the latent space from which it was decoded. (d, e, f) Distributions of mean FC, modularity, and FS-delta for the different sets of surrogate FCs generated by our model (respectively). All the distributions are significantly different (*p* < *0.001*, corrected with the Bonferroni correction). The effect sizes for these differences can be seen in the supplementary material (**Figure S1**).

To further characterize the different conditions, we computed the mean FC, modularity, and mean difference between the FC and the SC, referred to here as the Functional-Structural discrepancy (FS-delta). These metrics have been used previously in the literature [45], [46], [22], [47], [48] to differentiate between levels of consciousness, both physiological (e.g., sleep) and pathological (e.g., DoC). The metrics for all surrogate FCs are shown in **Figures 3d, 3e and 3f**.

The CT-DBS conditions, associated with recovery of consciousness, resulted in the lowest modularity, and highest mean FC and FS-delta, consistent with widespread cortical activation departing from the underlying SC. In contrast, both VL-DBS conditions exhibited FS-delta similar to that of the anesthesia condition and also showed reduced mean FC and increased modularity. Crucially, our analysis revealed that each DBS intervention target can be associated with a single direction in the latent space, corresponding to a particular direction related to the recovery of consciousness, which, at the same time, is a trajectory that increases the mean FC and FS-delta and decreases modularity [22].

### Wave perturbation most closely matches experimental transitions, validating the model against 3V VL-DBS dynamics

The 3V VL-DBS condition provided an appropriate reference point for our model, as it was reported to activate selectively the V2 region of the macaque cortex [34]. This cortical specificity enabled us to compare experimental data directly with our cortical model in an interpretable manner. Although CT-DBS was the most effective stimulation in restoring signatures of consciousness, it also induced widespread cortical and subcortical activation, making it unsuitable for validating our cortical model. Such broad effects would have made the validation process more complex, as they could arise from a myriad of interacting factors and obscure causal inference.

To validate our framework, we perturbed only V2 in the anesthesia model and tested whether the resulting dynamics resembled those observed under 3V VL-DBS. Perturbations were implemented using three approaches (i.e., noise, wave, and sync), each with an amplitude parameter. These allowed us to generate sets of trajectories in the latent space as a function of the perturbational amplitude.

Figure 4 shows a resulting trajectory for each perturbation type. By simulating each approach several times, we obtained distributions of latent-space distances between the dynamics induced by the perturbations and the centroids of the clouds of points relative to the experimental conditions. This enabled us to perform a direct comparison between the perturbational model and the augmented data.

As illustrated in Figure 4, bilateral sync perturbation of V2 failed to replicate the dynamics of 3V VL-DBS stimulation. Instead, it resulted in dynamics significantly closer to 3V CT-DBS than to 3V VL-DBS dynamics (*p < 0.05*), as indicated by the negative Cohen’s d value (*-0.47*). The wave perturbation outperformed the noise perturbation in specificity when reproducing 3V VL-DBS dynamics, with a Cohen’s d of *2.89* compared to *0.85* for noise between the distances to both low-voltage brain stimulation states. Furthermore, wave perturbation produced dynamics closer to 5V VL-DBS than noise perturbation (*p < 0.001*, Cohen’s d = *1.77*). This displacement reflects the directionality of the changes in dynamics.

**Figure 4.**
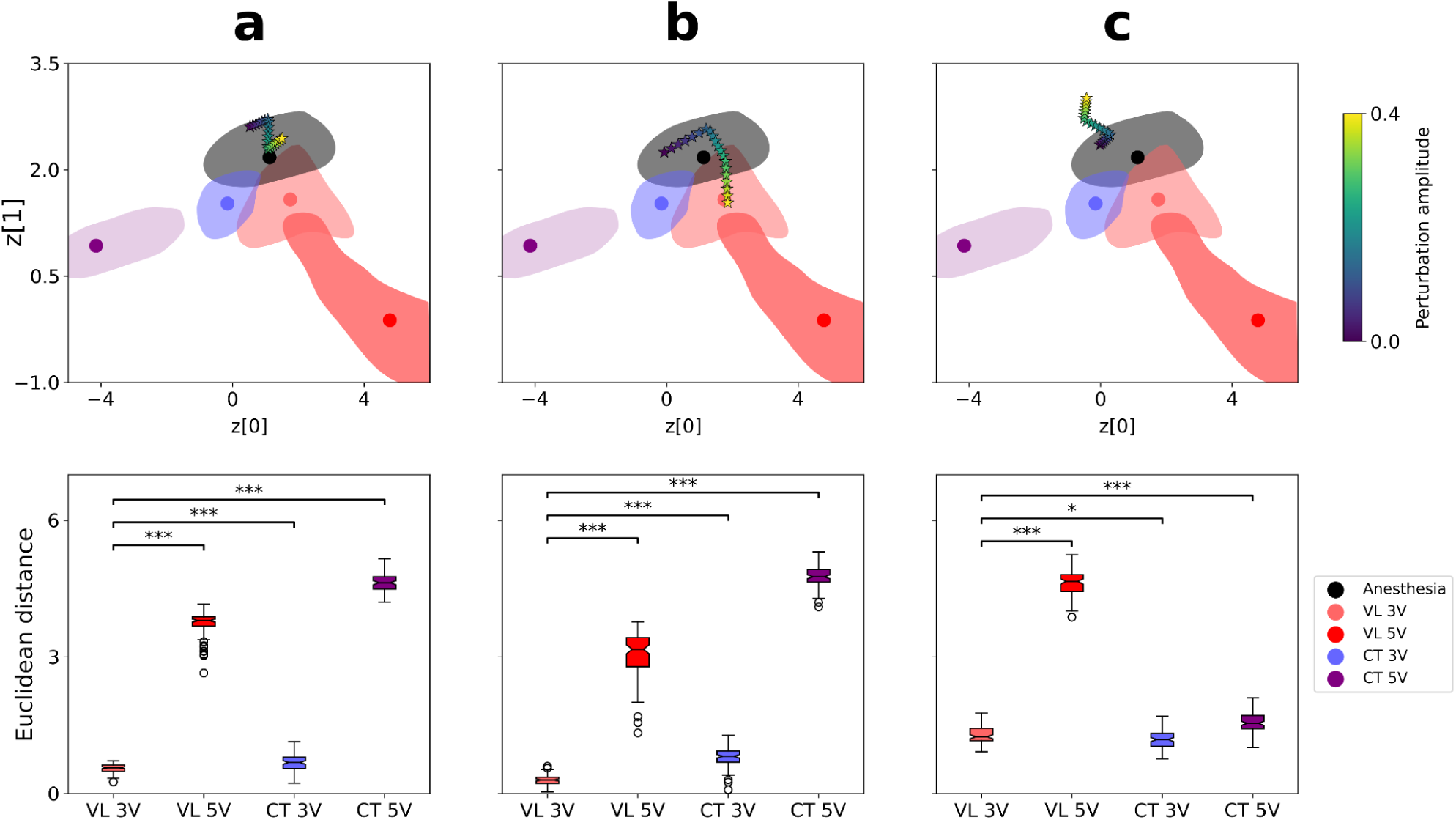
Matching experimental brain dynamics by perturbing the anesthesia model. (Top row) Trajectory of a perturbation of the model using different perturbation protocols (a: noise; b: wave; c: sync) targeting the secondary visual cortex (V2) bilaterally. The circular markers are the centroids of the point cloud for each condition. (Bottom row) Distribution of distances between the dynamics described by the perturbed model (21 amplitudes, 100 runs, 10% of lowest distances to each condition) and the centroid corresponding to each of the experimental conditions’ dynamics for each perturbational approach.

An exhaustive perturbation of all pairs of nodes to our anesthesia model further revealed limited flexibility of the noise and sync approaches (**Figure S2**). Noise perturbation consistently failed to reach CT-DBS dynamics, while sync perturbation was unable to reproduce VL-DBS states. Together, these findings support the continued use of wave perturbation in subsequent explorations of plausible perturbational targets, given its ability to specifically replicate 3V VL-DBS dynamics by targeting V2 bilaterally. In contrast, the noise and sync perturbational approaches were deprioritized due to their reduced specificity and flexibility. Notably, the observation that sync perturbations consistently drove the system toward CT-DBS dynamics, whereas noise perturbations displaced the dynamics only toward VL-DBS-like dynamics, underscores the importance of synchronized activity in the recovery of consciousness signatures. For these reasons, subsequent analyses were performed using the wave perturbation.

### Exploring the perturbative landscape of the anesthetized monkey

We continued by exhaustively perturbing every pair of homotopic nodes in the cortex, i.e. the same area in both hemispheres. We identified regions whose effects closely matched those in the experimental conditions. We also found that perturbations leading to 5V CT-DBS dynamics followed an ordered trajectory that passed through 3V CT-DBS dynamics, consistent with the expected progression observed in the real-world experiment. In the upper row of Figure 5 the trajectories are followed by different targets (shown at the top for each column) for different amplitude values. These trajectories also showed saturation: as the amplitude increased, the distance between the dynamics generated at an amplitude value and those at the next one decreased.

**Figure 5.**
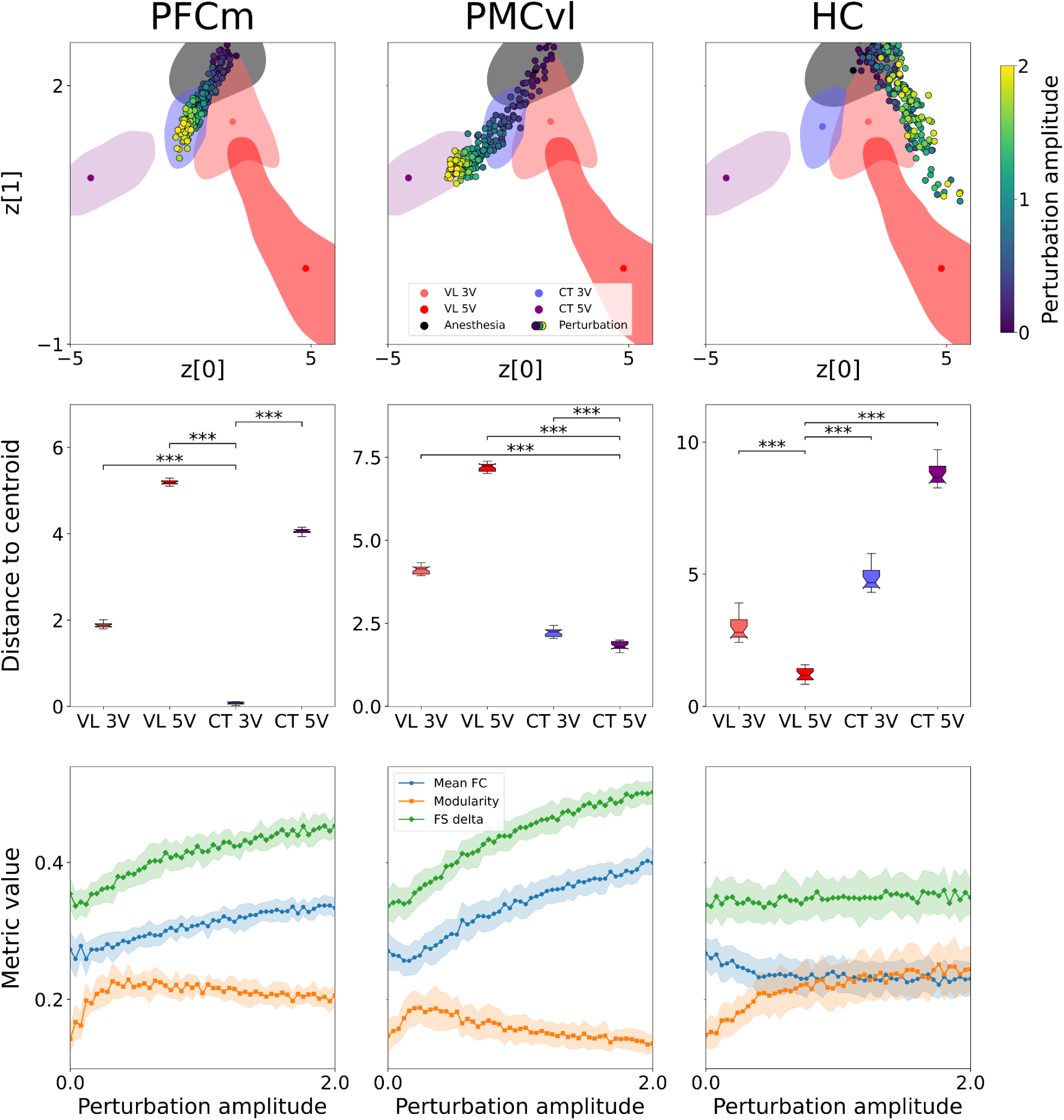
Perturbation of newly found cortical targets generate dynamics similar to those found during the thalamic stimulation in the experiment. Each column represents a different target for the perturbation: Medial Prefrontal Cortex (PFCm), Ventrolateral Premotor Cortex (PMCvl) and Hippocampus (HC). (Top row) Trajectories generated by perturbing the anesthesia model with different amplitudes using the wave perturbation for a single target. PFCm perturbation approaches 3V CT-DBS dynamics; PMCvl perturbation approaches 5V CT-DBS dynamics for high amplitudes and 3V CT-DBS for middle amplitudes; HC perturbation behaves as if it were just adding noise and gets close to the control conditions. (Middle row) For all the amplitudes used for perturbation, taking the top 10% of perturbations closest to the desired condition in the latent space, these boxplots show the distributions of distances to the centroid for each experimental condition. (Bottom row) The evolution of the different metrics (FS-delta in green with diamond markers being the upper line, mean FC in blue and round markers being the middle line, and modularity in orange and square markers being as the lowest line) as a function perturbation strength to the anesthesia model. All metrics are in the same value range as shown by the y-axis of the leftmost plot. The shaded area represents one standard deviation.

The dynamics obtained from perturbing the medial prefrontal cortex (PFCm) and the ventrolateral premotor cortex (PMCvl) encoded into the latent space led to the trajectories shown in Figure 5. These areas, under periodic perturbation, led the system’s dynamics to resemble those of CT-DBS stimulation. Other target areas found to produce a displacement toward the dynamics of the CT-DBS experimental condition were the temporal polar cortex (TCpol), the superior temporal cortex (TCs), the gustatory cortex (G), the centrolateral prefrontal cortex (PFCcl), the anterior cingulate cortex (CCa) and the medial premotor cortex (PMCm). Wave perturbation targeting other areas did not induce a change in dynamics in the direction of the CT-DBS dynamics. In particular, some parietal areas like the primary and secondary somatosensory cortices (S1 and S2), temporal areas like the primary auditory cortex (A1), frontal areas like the primary motor cortex (M1) and occipital areas like the primary visual cortex (V1) led to a displacement in the direction of VL-DBS dynamics under wave perturbation. For a more detailed view of the displacements generated by using different perturbation targets, **Table S1** has a summary of the distances of each perturbation to the different conditions (also picturized in **Figure S3**).

These perturbations were also specific, as the trajectories tended to go in a direction in a self-consistent manner. The distributions of distances from a perturbation to the centroid of each target condition, shown in the middle row of Figure 5, demonstrate this specificity. Notably, perturbation of these regions not only generated trajectories in the latent space but also brought the metrics, associated with the underlying dynamics, closer to those of the centroids they approached. Furthermore, perturbations of the PFCm and PMCvl induced dynamical regimes with signatures consistent with a departure from the anesthesia condition: increased mean FC [49], reduced modularity [50] and increase of FS-delta [51] (lower panels in Figure 5).

### Replication of perturbation-stimulation matching with data from a second monkey

To test whether our framework would hold up beyond the original dataset, we ran the same pipeline on data from a second macaque (monkey N). The available data was more limited, with only the anesthesia and the CT-DBS conditions were recorded, with also fewer overall trials. Because of this, the fitted models were less accurate and the VAE had to be trained without the control condition data. Not surprisingly, the latent space turned out to be less structured than for monkey T, with most of the variance captured along a dominant axis defined by the direction between the anesthesia state and the 5V CT-DBS state (**Figure S4**, left).

Despite this, the conditions separated clearly, and the ordering of both conditions and metrics showed the same general organization as in monkey T, the first animal (**Figure S4**, right). In the same fashion as with monkey T, perturbations to the anesthesia model of monkey N drove the dynamics to those of the stimulation conditions, with the same targets driving them toward CT-DBS states in both animals.

Further details of the replication are provided in Supplementary Section 4 and figures S4-S7.

### Choosing targets based on previous knowledge

Previous literature describes the macaque Global Neuronal Workspace (GNW) nodes network as a collection of higher-order prefrontal, cingular, and parietal regions that form a global workspace network [39]. It is mainly comprised of the posterior and anterior cingulate cortices (CCp and CCa), intraparietal cortex (PCip), the frontal eye field (FEF), the dorsolateral prefrontal and premotor cortices (PFCdl and PMCdl) and the prefrontal polar cortex (PFCpol). This network was validated in [34], [52]. For instance, in anesthetized macaques, the macaque Global Neuronal Workspace showed a striking decrease in functional correlation, with the posterior cingulate cortex and parietal cortex becoming disconnected from other bilateral GNW nodes. In contrast, awake states are characterized by strong intervoxel correlations within these prefrontal, parietal, and cingulate cortices [52]. Perturbation of the GNW network is hypothesized to drive a transition between brain states, such as from reduced consciousness (e.g., deep sleep or anesthesia) to higher consciousness. This would be achieved by influencing brain synchronization and reconfiguring the brain’s dynamic repertoire, specifically by increasing functional correlation and flexibility within these networks [16], [52].

For that reason, we decided to test how our model would react to a simultaneous perturbation of all of the GNW network’s nodes. In Figure 6a we can see that the dynamics under GNW perturbation move toward the CT for low perturbational amplitudes. However, with small increases to the amplitude, the trajectory changes and the brain state is found in an area of the latent space not inhabited by any of the experimental states. This perturbation saturates quickly and ends up at a specific location in the latent space. We hypothesize that this would be the area where the ‘awake’ brain state would lie. In Figure 6b we can see how the metrics evolve for the GNW perturbation. This perturbation leads to high mean FC and FS-delta and low modularity, which has been previously linked to recovery of consciousness.

**Figure 6.**
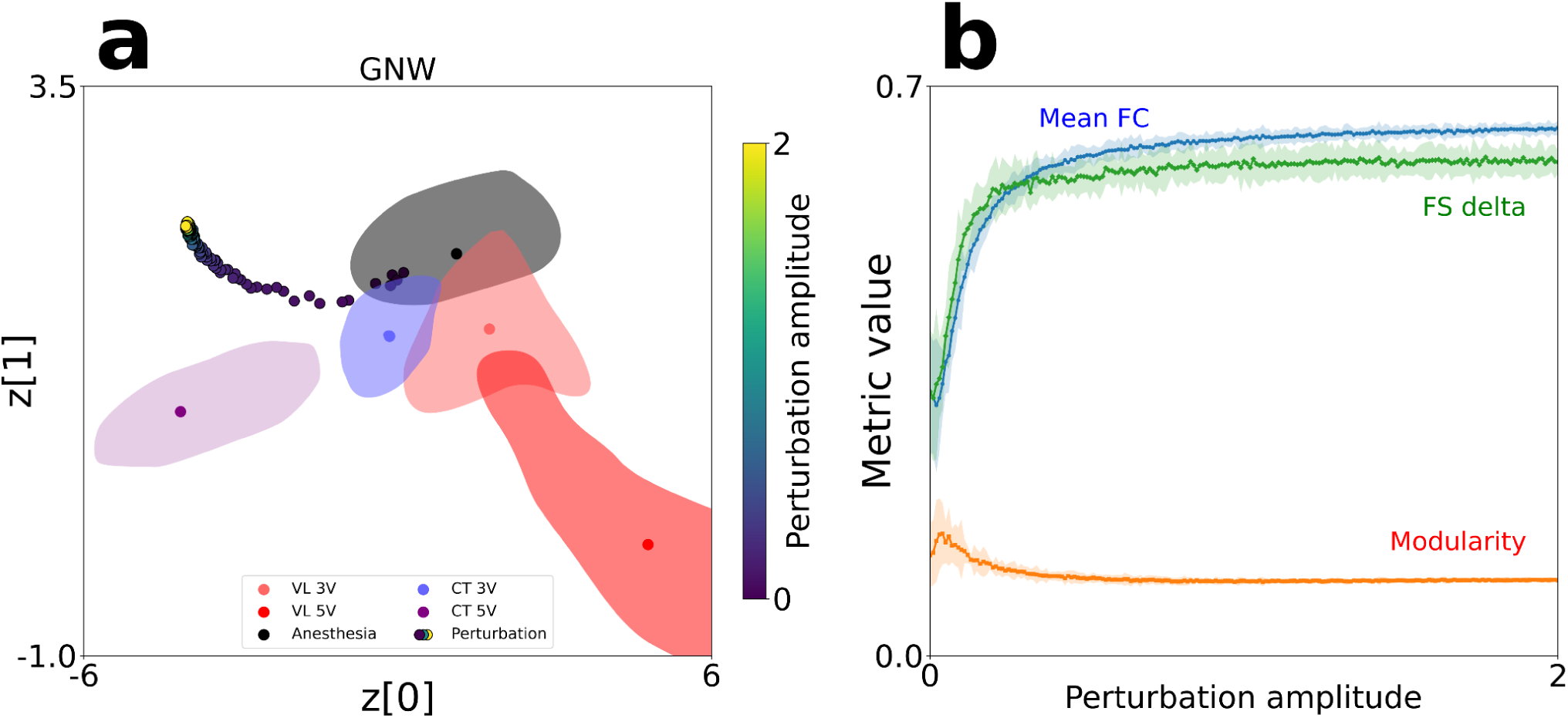
Perturbation of the macaque Global Neuronal Workspace network. (a) Trajectory generated by perturbing the anesthesia model with different strengths using the wave perturbation to all of the targets of the GNW network nodes. (b) The evolution of the different metrics (FS-delta in green and the upper line, mean FC in blue and middle line, and modularity in orange and as the lowest line) with perturbation strength in the dynamics generated by the model perturbation. All metrics are in the same value range as shown by the y-axis. The shaded area represents three standard deviations.

To further compare this perturbation with other states, we created a whole-brain model of an awake macaque (monkey B from the dataset) to obtain surrogate data to compare with the data from our perturbation. We also used empirical awake data from three different monkeys (monkeys B, J and Y from the dataset) to add a comparison to empirical data. To compare the different perturbational targets, we obtained the mean FC, modularity and FS-delta of the FC matrices obtained at the endpoint of each of the perturbations, and compared it with the values from said metrics of the surrogate data from the five models for different conditions for monkey T, the awake model for monkey B and the empirical awake data from monkeys B, J and Y.

## Discussion

We integrated two strategies to characterize and explore effective treatments for patients with disorders of consciousness (DoC): *in vivo* animal experimentation and *in silico* computational modeling. We used a unique dataset of macaque resting-state fMRI under propofol anesthesia, corresponding to an animal model of DoC, with and without intracranial electrical stimulation of different thalamic nuclei [34]. We then developed individualized whole-brain computational models tailored to the macaque’s fMRI data in various experimental conditions, enabling us to investigate the mechanisms underlying the restoration of consciousness. Importantly, we validated our model by capturing key aspects of *in vivo* results, by modeling stimulations in the cortical regions that were activated during the experiment. We continued by exhaustively exploring the full range of possible external interventions to assess whether different treatments could effectively restore consciousness. Afterward, we modeled perturbations to a network of nodes described as the global neuronal workspace network for macaques. And finally, we tested our results in a second monkey. The transitions between states were depicted as continuous trajectories in a low-dimensional space, identified in a data-driven manner by training variational autoencoders, enhancing the interpretability of the results. In summary, our findings demonstrate that whole-brain computational models can replicate the spatiotemporal characteristics observed in fMRI recordings during both the loss and recovery of consciousness, allowing us to investigate *in silico* perturbations and model potential interventions related to the restoration of consciousness.

A fundamental question is how dynamic interactions between brain regions, operating within a complex network of anatomical connections, lead to brain configurations associated with different states of consciousness. Computational modeling emerges as a suitable method to address this problem, as models can be used to contrast explanatory hypotheses concerning observed differences between brain states [53], [54], [55]. Among these differences, the correspondence between structural and functional connectivity has been consistently proposed as a marker of consciousness in humans, non-human primates and other animals like rats and mice [20], [34], [45], [46], [48], [56]. Importantly, this structure-function relation was also restored by CT-DBS in macaques [34], and was also correlated with the severity of disorders of consciousness. [46]. Recently, similar results were found using a multiscale approach that showed an increase in the coupling between brain activity and structural eigenmodes during pharmacological states in macaques [25] and in pharmacological and pathological loss of consciousness in humans [57]. In the case of mice and rats, it is also seen that the coupling between activity and structure is enhanced in the case of anesthesia [56] Our modeling efforts underscore the relevance of local dynamics to understand emergent patterns of structure-function coupling, as changes in the bifurcation parameter suffice to transition between states of consciousness, as indicated by changes in this and other signatures of consciousness.

Another relevant metric in this discussion is inter-areal coupling. Previous work has demonstrated that pharmacological and pathological reductions of consciousness in humans can be understood in terms of a reduction in global coupling strength, i.e., a decrease in the fiber conductivity of the structural traces [20]. Here, we showed that the anesthetized macaque presents the lowest coupling strength (G), while the states induced by CT-DBS stimulation, associated with a recovery of consciousness signatures, monotonically increase with the intensity of the stimulus, as is observed in humans during and after pharmacological intervention [20]. Conversely, VL-DBS stimulation did not present a significant increase of the coupling strength, supporting the association between the coupling strength inferred by the model and the states of consciousness. While anesthesia is not expected to modify the backbone of structural connections during the short induction time used in this experiment, an indirect effect on the effective connectivity can be expected based on how the drug interacts with local cell dynamics. In particular, propofol has been linked to increased neural inhibition, which could contribute to the functional decoupling of brain regions linked by long-range excitatory connections. In contrast to anesthesia, DoC patients may exhibit additional decoupling as a result of direct damage to the structural connectivity. Future modeling efforts could combine our approach with targeted attacks to the anatomical connectivity to better reproduce the behavior of DoC patients under invasive and non-invasive electrical stimulation.

Previously, Perl and colleagues [22] combined whole-brain modelling, data augmentation, and deep learning for dimensionality reduction, achieving a low-dimensional mapping capable of representing a continuum of states of reduced consciousness in a latent space. Interestingly, they found an orderly trajectory from wakefulness brain dynamics to those observed in patients with brain injuries, showing that one-dimensional manifolds can represent the brain state variability associated with a transition from wakefulness to deep unconsciousness. Here, we confirmed that a similar low dimensional representation was capable of capturing the changes in functional architecture measured during the restoration of consciousness in a single-macaque experiment.

Decoding signatures of consciousness from the latent space resulted in changes compatible with those reported in Perl et al., supporting the generalizability of our results across species and experimental conditions. Moreover, by suitably stimulating at a pre-defined region of interest we could reproduce the orderly transition toward consciousness represented in the manifold. This observation aligns with the previous results showing that restoration of consciousness, in terms of distributed cortical patterns [25] and in terms of behavioral measurements [34], can be triggered by selective stimulation of a specific subcortical region (CT) but not with others (VL). Moreover, we found that within each direction the stimulation amplitude is encoded as different orderly stops, i.e., strong stimulation is encoded far from the anesthetized state compared to weaker stimulation. Importantly, our model was capable of learning FC patterns that mapped into clearly separated latent space regions based on a single-subject model, while previous efforts in that direction were based on whole-brain models fitted to the average brain activity of several participants in certain brain states [22], [36], [58].

Once we showed that our model could reproduce the 3V VL-DBS transition, we exhaustively explored the full repertoire of transitions by applying the Wave protocol to perturb each pair of homotopic nodes in the macaque cortex. Crucially, we found that perturbations applied to specific temporal and prefrontal brain regions could restore the dynamics of the anesthetized macaque toward dynamical states associated with arousal and wakefulness. Conversely, the same perturbation applied to primary visual, auditory and somatosensory cortices (among others) was not prone to induce a restoration of the dynamics. Importantly, we were able to model which pair of brain regions is optimal for restoring brain dynamics, in agreement with a recovery of consciousness as reported in Tasserie et al. [34].

Studies in humans highlight that stimulation of the prefrontal cortex supports recovery of consciousness by enhancing connectivity within networks involved in awareness. This is particularly relevant for patients with DoC, where strengthening the fronto-parietal network has demonstrated promising results in improving both awareness and responsiveness [59], [60]. The prefrontal cortex, along with the parietal cortex, plays an important role in integrating external and internal information. Graph analysis of brain networks show how these areas are hubs of activity that play a role during complex cognitive processes [61]. The importance of these hubs for said characteristics, under the umbrella of consciousness, is supported by various studies using non-invasive brain stimulation techniques, such as transcranial direct current stimulation (tDCS). In these studies, stimulation of the dorsolateral prefrontal cortex (PFCdl) is proposed as a target for brain stimulation to assist with recovery from DoC [62], [24], [63], [64]. In this study we did not find PFCdl to be a target that displaces the dynamics away from those of the anesthesia condition. We believe that this happened due to the current limitations of our study: using only a cortical model and using macaque data, while the studies mentioned above are studies in humans. Nevertheless, the role of the prefrontal cortex in conscious experience and its role in supporting consciousness is still unclear [65] and current theories of consciousness argue about its role and the implication of its activity during different processes [66]. Some theories, like the GNW theory, give the PFC a high importance as a main hub for conscious experience [5], [67], while other theories, like IIT [3], [68] state that the PFC is not responsible for the conscious experience itself, but that its activity is related to the report of consciousness [69]. Instead, IIT proposes that the substrate of consciousness lies in a hotspot in the posterior cortex. We did not find any areas in the occipito-parietal cortex that, when perturbed, drove the dynamics away from the anesthesia state toward CT-DBS. The ongoing debate on the neural correlates of consciousness underscores its complexity as a distributed phenomenon dependent on both cortical as well as subcortical regions that were not modelled in this study.

Stimulation of primary sensory cortices typically yields localized sensory effects, such as the perception of phosphenes (perceived flashes of light or visual sensations without actual light stimulation) during occipital TMS or tactile sensations with somatosensory cortex stimulation [70]. Studies using tDCS in primary sensory areas show a modulation of excitability in the targeted regions and their associated networks with tDCS [71]. While these effects can provide insights into sensory processing and excitability, they do not seem to influence higher-order cognitive processes significantly. The differing effects of stimulating prefrontal areas versus primary sensory areas underscore the distinct functional roles of sensory and associative cortices. This highlights the critical importance of associative regions —whether it be the prefrontal cortex, posterior hotspots, or the prefrontal-parietal network— in developing therapeutic strategies to restore consciousness.

### Replicability

To evaluate the replicability of our findings, we used data from a second monkey to replicate the findings with the first one. Although not all of the experimental conditions were available in this second dataset, the framework revealed a consistent ordering of stimulation targets in the latent space, consistent with the findings for the first dataset. In both cases, perturbations of the same cortical areas produced similar changes in model dynamics toward those of the experimental conditions, despite inter-individual variability.

### Limitations

One of the major strengths of our study is the modelled and replication of *in vivo* experiments at single-macaque level, providing an individualized estimate for possible restoration of consciousness interventions. However, the dataset used in this work contains fMRI recording for one macaque in five experimental conditions and, consequently, our investigation is based on this single animal. Future works should extend this analysis for more animals and species. Nevertheless, it is limited by experimental difficulties that constrain the amount of available data. For the structural connectome data, a template derived from other studies, considered as a gold-standard in the field, was used in our work. We modelled the five experimental conditions using the same connectome with a two-step optimisation procedure that allowed us to have well-performing data-aligned models representing each FC: we consider that this limitation is not impacting in our results.

Our models and projections are based on cortical stimulations to induce restoration of consciousness. Nonetheless, subcortical structures are reported to be directly related to consciousness and stimulation in those regions are prone to restore consciousness states [31], [32], [33], [34]. In future work we will extend this approach by considering a finer parcellation and data from subcortical regions: we will explore which subcortical perturbations are prone to restore brain dynamics *in silico*.

The dataset for the second monkey lacked a control stimulation condition and had a lower number of trials for the available conditions, which hindered the fitting of our models and the expressiveness of the learned latent space by the VAE. We were able nonetheless to obtain perturbational targets that led the system to experiment dynamics, pointing toward the robustness of the framework used in the study. In order to fully explore the replicability of the study done with monkey T, a dataset containing all of the same conditions should be used.

### Future work

Future work on this line of inquiry should extend this framework beyond the single-subject, anesthetized data by incorporating datasets from multiple subjects under comparable stimulation conditions. Broadening the dataset will improve robustness and reproducibility, while studies in other species –in particular, humans– are essential to establish translational relevance. Incorporating data from patients with DoC would help capture the heterogeneity of etiology of said disorders, and enable the development of models tuned toward clinically meaningful predictions.

On the methodological side, incorporating subcortical regions should be a first key step in providing a more comprehensive view of the neuronal pathways underlying recovery of signatures of consciousness and allow for a broader exploration of possible stimulation targets. Finer parcellation would also improve the target search as it allows for higher spatial resolution.

Ultimately, advancing along both paths could lay the groundwork for computational frameworks that can reliably inform brain-stimulation-based treatments for patients with disorders of consciousness.

### Conclusions

Our computational framework applied to an animal model of unconsciousness successfully distinguished between brain-states arising from electrical stimulation and modeled the evolution of whole-brain dynamics in response to a localized perturbation. Importantly, these results were obtained using a data-driven approach. The latent space representation shown enabled a straightforward geometric interpretation of the stimulation data and the model perturbations, both in terms of matching between empirical stimulation states and *in silico* perturbational states, and in terms of relationship between different states.. The agreement between simulated and empirical latent space trajectories supports our approach and suggests its future applicability to new stimulation targets for the treatment of DoC patients. Our developments represent a step toward personalized treatments for individuals with disorders of consciousness, demonstrating that individualized whole-brain models can be developed and potentially used to identify targets for brain stimulation.

## Acknowledgments

**E.P.** is an FPI fellow funded by the Spanish “Ministerio de Ciencia, Innovación y Universidades” (MICIU/AEI/10.13039/501100011033) and “ESF investing in your future” under the grant PRE2020-0961.

**J.T., L.U.** and **B.J.** were supported by the Fondation pour la Recherche Médicale (FRM grant number ECO20160736100 to J.T.), Fondation Bettencourt Schueller, Fondation de France, Human Brain Project (Corticity project), Institut National de la Santé et de la Recherche Médicale, UVSQ, Commissariat à l’Energie Atomique, Collège de France.

**W.S.** was an FI fellow with the support of AGAUR, Generalitat de Catalunya and Fondo Social Europeo (2022 FI_B 00152) during this work.

**S.M.G.** was an FI fellow with the support of AGAUR, Generalitat de Catalunya and Fondo Social Europeo (2022 FI_B 00511) during this work.

**E.T.** was supported by grants FONDECYT Regular 1220995 (Chile), FONDECYT Exploración 13240170 (Chile) and CONICET-PIP 11220210100800CO (Argentina).

**M.L.K.** is supported by the Centre for Eudaimonia and Human Flourishing (funded by the Pettit and Carlsberg Foundations) and Center for Music in the Brain (funded by the Danish National Research Foundation, DNRF117).

**A.I.L.** is supported by St John’s College, Cambridge; and a Wellcome Early Career Award (grant number 226924/Z/23/Z).

**G.D.** was supported by the Grant PID2022-136216NB-I00 funded by MICIU/AEI/10.13039/501100011033 and by “ERDF A way of making Europe,” ERDF, EU.

**G.D.** is also supported by AGAUR research support grant (2021 SGR 00917) funded by the Department of Research and Universities of the Generalitat of Catalunya.

**Y.S.P.** is also supported by the European Union’s Horizon 2020 research and innovation program under the Marie Sklodowska-Curie grant 896354.

**G.D.** and **Y.S.P.** were supported by the project NEurological MEchanismS of Injury, and Sleep-like cellular dynamics (NEMESIS) (ref. 101071900) funded by the EU ERC Synergy Horizon Europe.

## Author contributions

- Conceptualization: E.P., Y.S.P., G.D., B.J.
- Methodology: E.P., C.G., S.M.G., G.D., Y.S.P.
- Software: E.P., W.S., Y.S.P.
- Simulations: E.P.
- Formal Analysis: E.P., Y.S.P.
- Data Curation: J.T., L.U., C.G., A.G., B.J.
- Visualization: E.P., Y.S.P.
- Writing – Original Draft: E.P., Y.S.P.
- Writing – Review & Editing: All authors
- Supervision: G.D., B.J., Y.S.P.
- Funding Acquisition: G.D.

## Methods

All information regarding the animal procedures, DBS implantation, anesthetic procedures, data acquisition and related methodology is available in [34]. Readers are referred to that publication for more in-depth methodological details. The following paragraphs serve as a summary.

### Data acquisition summary

#### Animals and surgical procedures

Research was conducted on five rhesus macaque monkeys (*Macaca mulatta*) following the European convention for animal care (86-406) and the National Institutes of Health’s Guide for the Care and Use of Laboratory Animals. The studies were approved by the Institutional Ethical Committee (CETEA protocols #12-086 and #16-040). For the awake resting-state experiments, monkeys B, J, and Y were implanted with an MR-compatible headpost under general anesthesia. Monkeys N and T were implanted with a clinical DBS electrode (Medtronic, Minneapolis, MN, USA, lead model 3389) with four active contacts. The electrode was implanted in the right CM thalamus, and the lead localization was verified postsurgically using a reconstruction method based on in vivo brain imaging and a postmortem histology study in one of the implanted monkeys.

#### fMRI data acquisition

Monkeys were scanned on a 3-T horizontal scanner (Siemens PrismaFit, Erlanger, Germany) with a customized eight-channel phased-array surface coil (KU Leuven, Belgium). For the DBS and resting-state experiments, echo planar imaging (EPI) was used, with a TR of 1250 ms, an echo time (TE) of 14.20ms, 1.25-mm isotropic voxel size, and 325 and 500 brain volumes per run, respectively.

For the awake protocol, animals were trained to sit in a sphinx position in a primate chair with their head fixed, without any task, and the eye position was monitored. The anesthesia for the DBS experiments was induced with an intramuscular injection of ketamine (10 mg/kg; Virbac, France) and dexmedetomidine (20 μg/kg; Ovion Pharma, USA) and maintained with a target-controlled infusion (TCI) (Alaris PK Syringe pump, CareFusion, CA, USA) of propofol (Panpharma Fresenius Kabi, France) using the “Paedfusor” pharmacokinetic model (monkey T: TCI, 4.6 to 4.8 μg/ml; monkey N: TCI, 4.0 to 4.2 μg/ml). A muscle-blocking agent (cisatracurium, 0.15 mg/kg, bolus i.v., followed by continuous intravenous infusion at a rate of 0.18 mg/kg per hour; GlaxoSmithKline, France) was used during all anesthesia fMRI sessions to avoid artifacts.

#### Electrical stimulation for the DBS experiments

The DBS electrode was connected to an external stimulator (DS8000, World Precision Instruments, USA), and the signal parameters were set to a monopolar signal at 130.208 Hz. The width of the pulse was different between the monkeys (320 μs was used for monkey N; 140μs was used for monkey T). The absolute voltage amplitude was set to 3V or 5V. The DBS fMRI data was acquired under anesthesia with 3V or 5V DBS on either the Central Thalamic nuclei (CT) or the Ventrolateral Thalamic nuclei (VL).

#### fMRI preprocessing

fMRI images were preprocessed using Pypreclin (Python preclinical pipeline) [72]. Functional images were corrected for slice timing and B_0_ inhomogeneities, reoriented, realigned and resampled (1.0 mm isotropic), masked, coregistered to the MNI macaque brain template [73], and smoothed (3.0 mm Gaussian kernel) using FSL (https://fsl.fmrib.ox.ac.uk/fsl/) [74] and custom Python code [39]. Regression of the movement parameters resulting from rigid-body correction was removed, along with the global signal, to rule out confounding effects from physiological changes. Voxel time series were filtered using a high-pass (0.0025 Hz cutoff) and low-pass (0.05 Hz cutoff) filters and with a zero-phase fast-Fourier (FFT) notch filter (0.03 Hz) to remove an artifactual pure frequency present in all sessions. The variance of each time series was also normalized; making it so that covariance matrices correspond to correlation matrices.

### Anatomical connectivity

The anatomical connectome used was a fully weighted whole-cortex macaque structural connectivity matrix (connectome) derived by combining the information from fiber-tracing and tractography [75]. The connectome is publicly available: https://zenodo.org/record/1471588#.X44C6dAzY2x. The tractography algorithm was optimized to best reproduce the weighted but partial-cortex tracer connectome from Markov et al. [76], before estimating the whole-cortex connectome weights. For details see Shen et al. [75].

### Whole-brain models

The whole-brain model used in this study is the denominated Hopf model [77]. It consists of a network of nonlinear oscillators (Stuart-Landau oscillators), each representing one brain region, coupled by the anatomical connectivity. To implement the model, two main assumptions are made: (i) the dynamics of macroscopic neural masses can range from fully synchronous to a stable asynchronous state driven by random fluctuations; (ii) fMRI can capture the dynamics from both regimes with sufficient consistency and reliability to be modeled by the equations. Each oscillator represents the dynamics of one of the 82 regions of the parcellation. The Hopf bifurcation changes the qualitative nature of the solutions from a stable fixed point to a limit cycle, allowing the model to represent the emergence of self-sustained oscillations. The dynamics of brain region *j* were modeled by the following equations:

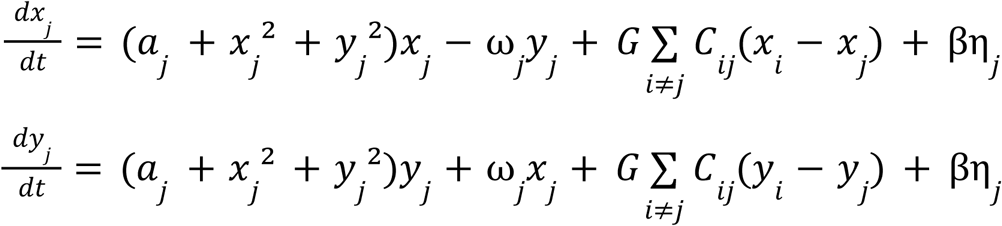

In the equations, ω𝑗 is the intrinsic oscillation frequency of node *j*. This intrinsic oscillation frequency is the highest-powered frequency from the BOLD signal of each area. The parameter 𝑎_𝑗_ is known as the bifurcation parameter, and the dynamics of the system change from a stable fixed point to a limit cycle depending on the value of this parameter. The coupling comes into play in the third term of the equations. Nodes *i* and *j* are coupled by *C_ij_* (the *i,j* entry of the SC matrix). β_j_ is fixed at 0.04 and represents the scaling factor of the additive white Gaussian noise (η_j_) for each node. The parameter *G* is the global coupling factor and scales the SC equally for all nodes. The empirical SC matrix was globally scaled to a maximum of 0.2 (weak coupling assumption). The equations were integrated using the Euler-Maruyama algorithm with a time-step of 0.01 seconds to simulate the empirical fMRI signals. In this model, when *a* is close to the bifurcation, the additive white Gaussian noise gives rise to complex dynamics as the system continuously switches between both sides of the bifurcation.

The BOLD signal for each node is obtained from the integrated x_j_ value. Afterward, a bandpass filter is used to bandpass the signal in order to make it comparable to the empirical one, using the same cutoff frequencies than the ones used in the experimental study (0.0025 Hz lower cutoff frequency, and 0.05 higher cutoff frequency, Butterworth filter of order 2). No notch filter is necessary in the case of this data due to the artifactual pure frequency not being present in the simulated data.

The implementation of this model can be found here: https://github.com/decolab/restoring-signatures-individual-WBM

### Model fitting - Parameter optimization procedure

The fitting procedure consisted of two steps: first, the global coupling parameter was optimized, followed by the fitting of the bifurcation parameters for each pair of brain areas. The empirical observable used for model fitting was the trial-averaged single-subject functional connectivity (FC), also called the zero-lag normalized covariance matrix. A model was created for each experimental condition of monkey T (anesthesia, 3V CT stimulation, 5V CT stimulation, 3V VL stimulation and 5V VL stimulation).

The above described whole-brain model was used to simulate regional fMRI signals. The first step toward fitting the model is exhaustively exploring the parameter space for the coupling parameter (G), using a homogenous condition for the model with the bifurcation parameter of every region close to the bifurcation (a ≈ 0, a = -0.02), which according to previous literature leads to the best fit in a homogenous model [77]. The metric used to fit the data was the structural similarity index (SSIM) [78] as implemented by the scikit-image package under the SciPy package [79] in the Python programming language. This metric is what we will use to determine the goodness of fit (GoF). This metric had been used previously in the context of model fitting for data augmentation with human data under different experimental conditions [40], and was used to provide a more nuanced fitting than using only Pearson’s correlation while still allowing the use of a single metric. From the exhaustive exploration, the *G* value that maximizes the SSIM is selected.

Afterward, a fitting of the *‘a’* parameters is done by the application of a genetic algorithm following previous work [43]. In this case, instead of lowering the dimensionality of the parameter space for the bifurcation parameters by using resting-state networks or other networks, the dimensionality was reduced from the 82 different areas to 41 parameters. The assumption here is that homotopic areas in both hemispheres would have a similar enough cytoarchitecture to have similar local dynamics. In a work by Shamir and colleagues, they report that the cytoarchitecture of homotopic areas is highly symmetric [45]. The drawback of this assumption is that it cannot represent asymmetries (e.g., interhemispheric) which have been reported by literature [80]. Then, the individuals of each generation would have what is called a genome of 41 parameters, which is used to run the Hopf model and obtain the SSIM between the model and the empirical data. Each generation, composed of 20 individuals, would then be ranked by the GoF and the two highest models go on intact to the next generation (elite selection). The rest of the 18 individuals are made up of children of the individuals from the previous generation. Parents are selected with different probabilities depending on their GoF (better GoF means more probability to be picked), and their genomes are mixed to create a new individual (crossover). The genomes of these individuals have the chance to mutate, changing the value of a random parameter following a normal probability distribution centered on the respective value (mutation). This process goes on for 200 generations or until the highest GoF has not varied more than a given value (we used 0.00001) for the last 10 generations (i.e. if it has converged). This process is run 100 times and the parameters of the model that outperforms the rest are chosen as the ones for the model to run the data augmentation.

### Data augmentation

The best performing fitted model was used to generate surrogate data for later training of the VAE. 5000 FC matrices were generated of the anesthesia, 5V CT-DBS, and 5V VL-DBS conditions of monkey T for training the VAE. Additionally, another 500 FC of all conditions were generated for analysis and visualization purposes, in order for them not to be from the training or validation sets, i.e. so that they are unknown to the VAE model.

### Model perturbation - In silico stimulation of the brain

Perturbation to the models was done following the procedures in Sanz Perl et al. (2022) [36]. Two main procedures were used to perturb an area: applying an extra oscillatory term (wave stimulation), and shifting the value of the bifurcation parameter of said area (called noise stimulation if decreasing the value, or sync stimulation if increasing the value). The wave stimulation is given by an additive term given by F_0_cos (ω_0_t). F_0_ is the forcing amplitude, and it varies from 0 to 2 in steps of 0.1 in order to parametrize the perturbation as a function of the strength. ω_0_ is the natural frequency of the node that is being stimulated. Each simulation of the perturbation was made with 20 sub-simulations to obtain a mean FC.

For figure 4, interpolated versions of the trajectories described by the perturbations were created for visualization purposes. The distances shown in boxplots are calculated as follows: For a given perturbation target the perturbation is parameterized by the perturbation amplitude from 0 to 2 in 21 steps. The perturbation is simulated 100 times. The distance between every strength value in every simulation and all the centroids of the clouds of points is calculated and then, the distances of the perturbations with the 10% lowest distance to each condition was taken as the distance between that perturbational target and said condition.

For figure 5, to calculate the metrics (mean FC, modularity and mean difference between FC and SC, namely Functional-Structural discrepancy or FS-delta for short) parameterized by the perturbation strength, a different amount of simulations was used: 20 simulations were performed, with 5 sub-simulations per simulation (to obtain the FC) and the strength was parameterized from 0 to 2 in 51 points.

For figure 6, we used a parameterization of 201 points between 0 and 2, with 10 simulations with 10 sub-simulations. In the case of the metrics for the empirical awake data, we did not group the trials together, calculating the mean FC, modularity and FS-delta of each trial separately.

### Variational Autoencoder (VAE)

The VAE implementation used follows that of Sanz Perl and colleagues (2022) [22]. The data encoded into the latent space are the FC matrices, which for training purposes are then decoded and the reconstruction error is minimized. The VAE maps inputs to probability distributions in the latent space, which are regularized during this training process. This allows the latent space to become meaningful, meaning that coordinates not used by empirical data can still be decoded and produce interpretable results.

The architecture of the VAE follows the one used in [22]. Briefly, they consist of three parts: the encoder networks, a middle variational layer and the decoder network. The encoder performs a dimensionality reduction from the data space to a latent space. The decoder then reconstructs the input from the information in the latent space.

The training is done with the generated matrices from monkey T under the conditions of anesthesia, 5V CT-DBS and 5V VL-DBS. The training procedure consisted of batches of 256 samples and 5 epochs. Later, generated FCs from all states are encoded for visualization in the latent space. The decoder is also used for decoding an array of points of the latent space for study of the different metrics that characterize it.

The grid used to decode the latent space was a 20×20 grid of equally spaced points in the range seen in Figure 3c.

### Metrics used

Outside of the latent space, three metrics are used to characterize the dynamics of each model (both unperturbed and perturbed): mean FC, modularity and FS-delta. The mean FC is calculated as the average of all elements of the FC matrix. Modularity is computed using the NetworkX python package [81]. Specifically, the FC matrix is treated as an adjacency matrix of a weighted graph, and communities are detected using the Louvain algorithm. Then, modularity is calculated from the obtained communities. The final modularity value is obtained from the average value after 10 runs from this process, in order to obtain some stability from the community detection. Finally, the Functional-Structural Discrepancy (FS-delta for short) is calculated by subtracting the normalized SC from the normalized FC and then calculating the mean of the values of the resulting matrix. The choice in naming for this metric was so to avoid confusion with the euclidean distance used to characterize states in the latent space.

Within the latent space, the metric used to characterize and compare states was the euclidean distance.

### Statistical analyses

Statistical significance was obtained using the Kruskal-Wallis test, as implemented in the SciPy computing library [79]. Bonferroni correction for multiple comparison was applied when necessary. Differences were considered significant (*) when the corrected p-value was below 0.05, very significant (**) when it was below 0.01 and highly significant (***) when below 0.001. Effect sizes were calculated using Cohen’s d.

