## Supplementary Materials for "Restoring signatures of consciousness by thalamic stimulation in a whole-brain model of an anesthetized nonhuman primate"

### *Supplementary information*

*Section 1. Effect sizes calculated between distributions of different metrics from different conditions (monkey T augmented data)*

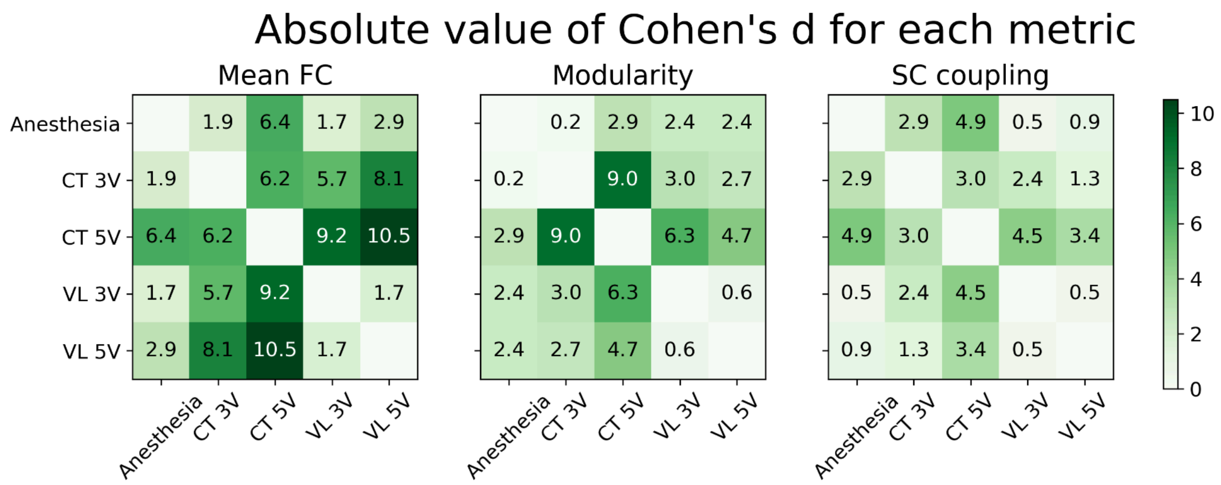


**Figure S1. Effect sizes.** For each different metric of dynamics, we calculated the effect sizes between the distributions given by the different simulations of each condition. This figure complements the information from figure 3 panels d, e and f.

*Section 2. Flexibility of perturbation types*

In **Figure S2** the trajectories in the latent space obtained from perturbation to all 41 pairs of nodes with each perturbation type is represented. Each of the 41 perturbational targets is represented by 201 points for each different perturbational amplitude from 0 to 2.

Noise and sync perturbation lead the dynamics dominantly in one direction, while wave perturbation shows the ability to move in both directions depending on the target of said stimulation, even if with less displacement per unit of perturbational amplitude than, for example, the sync perturbation. This capacity to move in different directions is the flexibility that allows this approach to simulate the dynamical effects of both DBS stimulation targets.

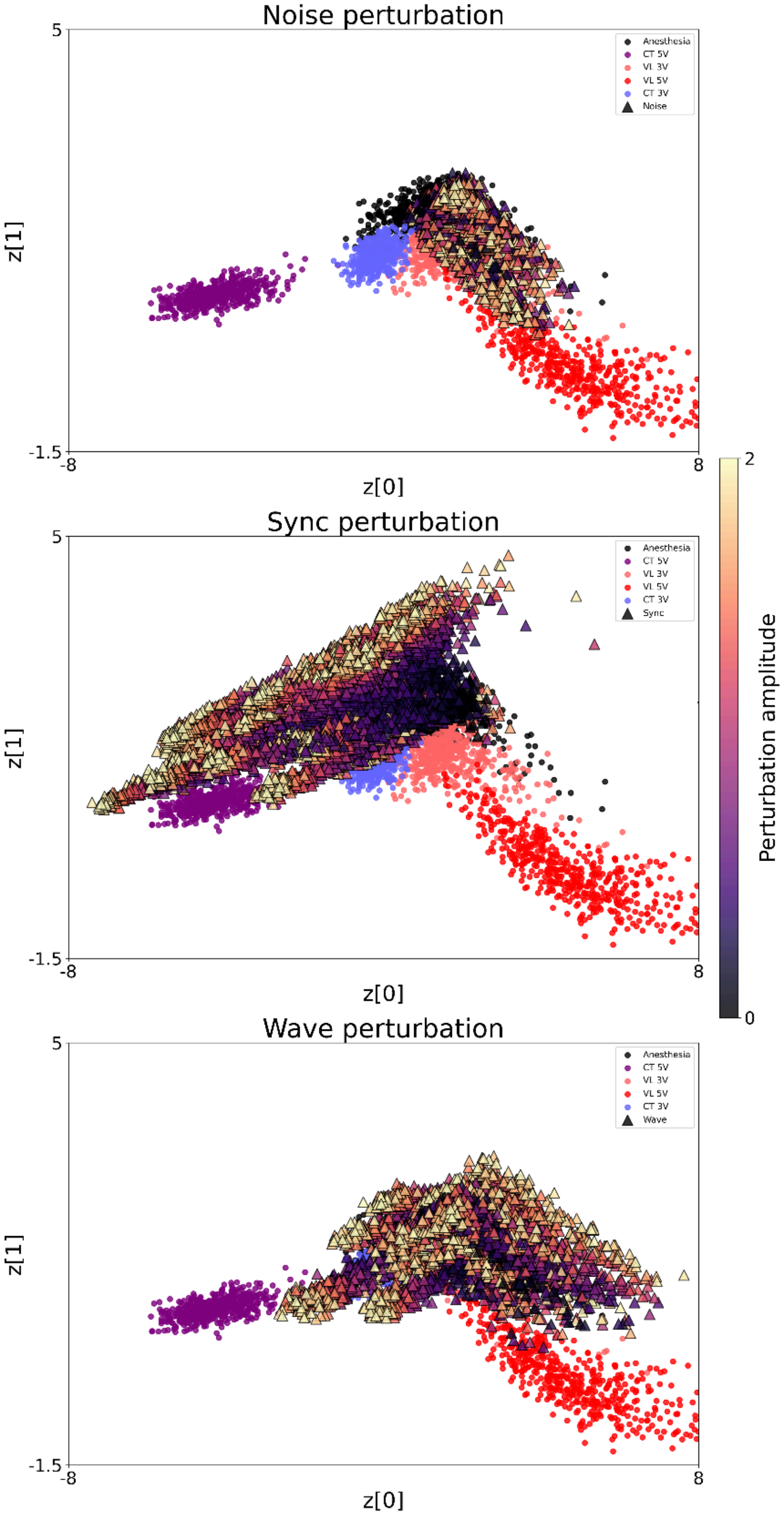


**Figure S2. Perturbation type flexibility.** For each different perturbation type (noise, sync and wave) we perturbed each pair of homotopic nodes (41 pairs in total) with varying amplitude (0 to 2 in 201 steps).

*Section 3. Distances of different stimulations to the centroids of the experimental conditions*

**Table S1. Mean distance in the latent space of the perturbational dynamics to each of the experimental dynamics clusters' centroids.** This table shows the mean ± standard deviation for each perturbational target. The results shown represent the mean and standard deviation of the 10% of strength values that minimize the distance between the dynamics obtained and the corresponding centroid. The anesthesia condition is not in the table as it is the starting condition for the perturbation.

| | Distance to … condition | | | |
|---|---|---|---|---|
| **Area** | **3V CT-DBS** | **5V CT-DBS** | **3V VL-DBS** | **5V VL-DBS** |
| Tempolar polar (TCpol) | 0.38 ± 0.08 | 4.27 ± 0.1 | 0.62 ± 0.04 | 3.87 ± 0.1 |
| Superior temporal cortex (TCs) | 0.55 ± 0.04 | 3.29 ± 0.1 | 0.58 ± 0.1 | 3.75 ± 0.2 |
| Amygdala (Amyg) | 1.66 ± 0.2 | 5.63 ± 0.2 | 0.70 ± 0.04 | 2.96 ± 0.2 |
| Orbitoinferior prefrontal cortex (PFCoi) | 1.70 ± 0.2 | 5.65 ± 0.2 | 0.72 ± 0.07 | 2.86 ± 0.4 |
| Anterior insula (Ia) | 2.00 ± 0.2 | 5.98 ± 0.2 | 0.78 ± 0.06 | 2.95 ± 0.3 |
| Orbitomedial prefrontal cortex (PFCom) | 1.51 ± 0.08 | 5.49 ± 0.08 | 0.57 ± 0.05 | 3.63 ± 0.09 |
| Central temporal cortex (TCc) | 0.97 ± 0.03 | 4.29 ± 0.1 | 0.76 ± 0.09 | 3.91 ± 0.2 |
| Orbitolateral prefrontal cortex (PFCol) | 1.82 ± 0.09 | 5.78 ± 0.08 | 0.70 ± 0.07 | 2.99 ± 0.5 |
| Inferior temporal (TCi) | 1.46 ± 0.1 | 5.43 ± 0.1 | 0.51 ± 0.07 | 2.84 ± 0.3 |
| Parahippocampal cortex (PHC) | 2.17 ± 0.2 | 6.17 ± 0.2 | 0.76 ± 0.1 | 1.29 ± 0.1 |
| Gustatory cortex (G) | 0.13 ± 0.07 | 2.76 ± 0.1 | 0.79 ± 0.08 | 4.08 ± 0.2 |
| Ventrolateral premotor cortex (PMCvl) | 0.16 ± 0.07 | 1.83 ± 0.1 | 0.83 ± 0.1 | 4.18 ± 0.2 |
| Anterior visual area (ventral) (VACv) | 1.77 ± 0.1 | 5.78 ± 0.1 | 0.36 ± 0.1 | 1.54 ± 0.3 |
| Posterior insula (Ip) | 1.84 ± 0.09 | 5.82 ± 0.1 | 0.63 ± 0.08 | 2.81 ± 0.3 |
| Prefrontal polar cortex (PFCpol) | 1.09 ± 0.07 | 5.06 ± 0.06 | 0.69 ± 0.04 | 3.83 ± 0.2 |
| Hippocampus (HC) | 2.02 ± 0.2 | 6.01 ± 0.2 | 0.71 ± 0.06 | 1.21 ± 0.2 |
| Subgenual cingulate cortex (CCs) | 1.84 ± 0.07 | 5.81 ± 0.07 | 0.70 ± 0.04 | 3.00 ± 0.2 |
| Ventrolateral prefrontal cortex (PFCvl) | 0.95 ± 0.2 | 4.91 ± 0.2 | 0.64 ± 0.05 | 3.26 ± 0.5 |
| Visual area 2 (V2) | 1.81 ± 0.1 | 5.82 ± 0.1 | 0.27 ± 0.09 | 1.95 ± 0.2 |
| Medial prefrontal cortex (PFCm) | 0.08 ± 0.03 | 3.80 ± 0.1 | 0.76 ± 0.07 | 4.07 ± 0.1 |
| Ventral temporal cortex (TCv) | 1.51 ± 0.07 | 5.48 ± 0.09 | 0.46 ± 0.07 | 2.95 ± 0.3 |
| Anterior visual area (dorsal) (VACd) | 2.69 ± 0.7 | 6.71 ± 0.7 | 1.07 ± 0.5 | 1.24 ± 0.2 |
| Visual area 1 (V1) | 1.77 ± 0.09 | 5.79 ± 0.1 | 0.21 ± 0.07 | 1.82 ± 0.2 |
| Centrolateral prefrontal cortex (PFCcl) | 0.13 ± 0.06 | 4.09 ± 0.08 | 0.69 ± 0.07 | 3.85 ± 0.2 |
| Secondary auditory cortex (A2) | 0.69 ± 0.1 | 4.67 ± 0.1 | 0.17 ± 0.04 | 3.11 ± 0.4 |

| | | | | |
|---|---|---|---|---|
| Retrosplenial cingulate cortex (CCr) | 1.67 ± 0.1 | 5.67 ± 0.1 | 0.55 ± 0.05 | 2.77 ± 0.2 |
| Posterior cingulate cortex (CCp) | 1.60 ± 0.09 | 5.49 ± 0.1 | 0.77 ± 0.05 | 3.52 ± 0.3 |
| Anterior cingulate cortex (CCa) | 0.23 ± 0.07 | 1.91 ± 0.2 | 0.82 ± 0.1 | 4.09 ± 0.3 |
| Secondary somatosensory cortex (S2) | 0.47 ± 0.05 | 3.85 ± 0.1 | 0.26 ± 0.09 | 3.25 ± 0.3 |
| Primary somatosensory cortex (S1) | 1.97 ± 0.2 | 5.97 ± 0.2 | 0.50 ± 0.06 | 1.39 ± 0.3 |
| Primary auditory cortex (A1) | 0.93 ± 0.1 | 4.97 ± 0.1 | 0.28 ± 0.05 | 3.13 ± 0.6 |
| Primary motor cortex (M1) | 1.63 ± 0.1 | 5.65 ± 0.1 | 0.28 ± 0.08 | 2.24 ± 0.4 |
| Inferior parietal cortex (PCi) | 3.04 ± 0.3 | 7.04 ± 0.3 | 1.34 ± 0.3 | 1.46 ± 0.2 |
| Medial parietal cortex (PCm) | 1.88 ± 0.2 | 5.84 ± 0.2 | 0.82 ± 0.06 | 1.93 ± 0.1 |
| Dorsomedial prefrontal cortex (PFCdm) | 1.82 ± 0.1 | 5.81 ± 0.1 | 0.65 ± 0.06 | 2.75 ± 0.2 |
| Intraparietal cortex (PCip) | 2.19 ± 0.3 | 6.19 ± 0.3 | 0.81 ± 0.09 | 1.88 ± 0.2 |
| Superior parietal cortex (PCs) | 2.02 ± 0.2 | 6.01 ± 0.2 | 0.76 ± 0.07 | 1.68 ± 0.2 |
| Frontal eye field (FEF) | 1.45 ± 0.1 | 5.44 ± 0.1 | 0.60 ± 0.06 | 2.64 ± 0.3 |
| Dorsolateral prefrontal cortex (PFCdl) | 1.28 ± 0.2 | 5.25 ± 0.2 | 0.63 ± 0.07 | 2.56 ± 0.5 |
| Medial premotor cortex (PMCm) | 0.45 ± 0.1 | 4.38 ± 0.1 | 0.63 ± 0.04 | 3.68 ± 0.5 |
| Dorsolateral premotor cortex (PMCdl) | 1.46 ± 0.1 | 5.44 ± 0.1 | 0.64 ± 0.04 | 3.30 ± 0.2 |

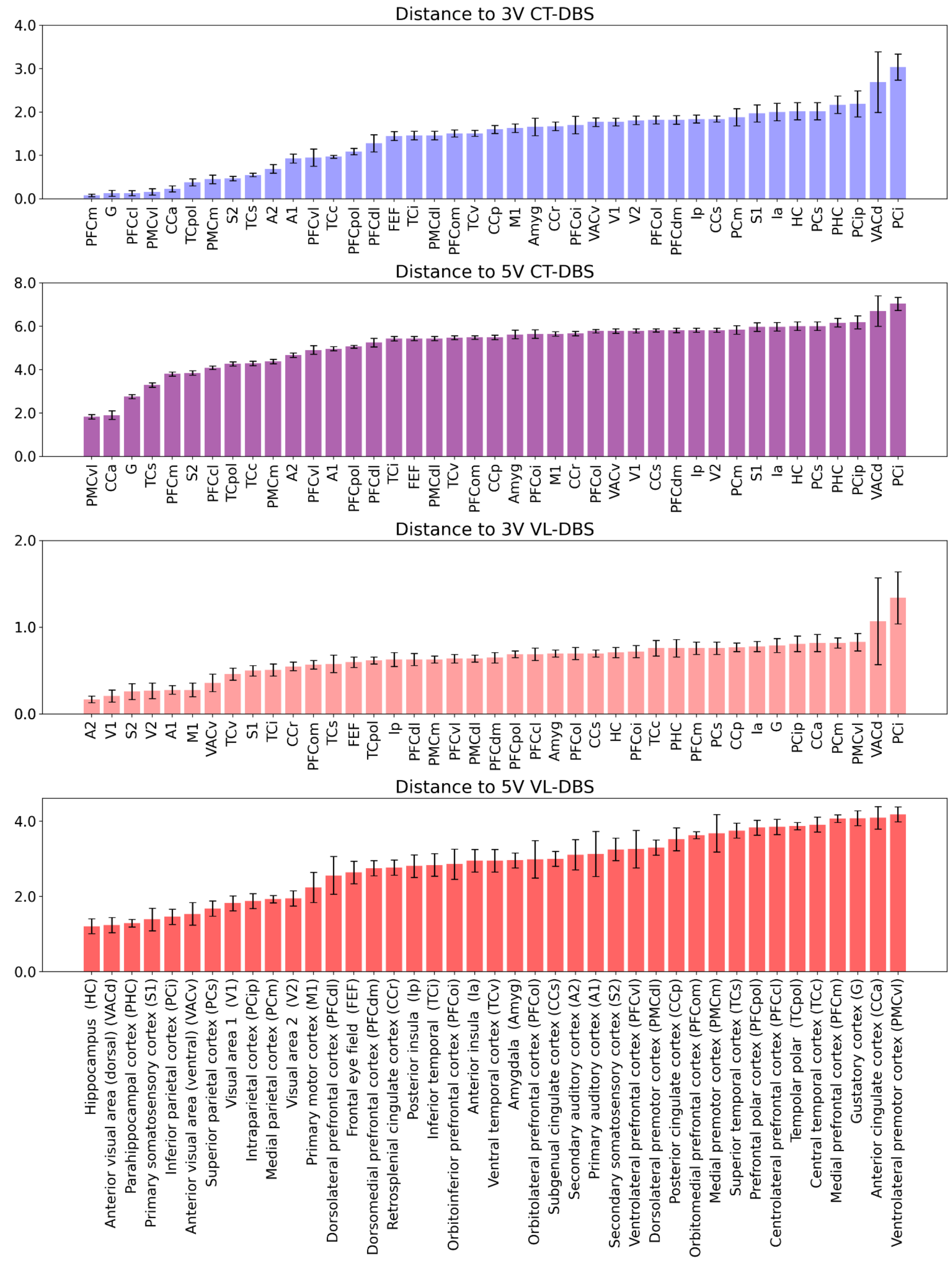


**Figure S3. Mean distance in the latent space of the perturbational dynamics to each of the experimental dynamics clusters' centroids.** Depiction of the results from **Table S1** per target condition. Ordered from lowest to highest per condition.

*Section 4. Replication in a second monkey (Monkey N)*

To validate our framework, we replicated the analysis using data from a second macaque, monkey N. The dataset of monkey N included recordings under anesthesia and under CT-DBS stimulation at 3V and 5V. The same modeling pipeline as for monkey T was used.

A VAE was trained using the three available conditions. The latent space obtained is shown in the left panel of **Figure S4**. The right panels display the mean FC, modularity, and FS-delta for each condition, which maintain the same ordering as in the case of monkey T, indicating robustness across subjects. The differences between conditions for every metric were highly significant after correction (*p < 0.001*). The effect sizes for the comparisons are reported in **Figure S5**. As with monkey T, the differences between anesthesia and 5V CT-DBS have the largest effect size. The overall model fit was lower than for monkey T due to fewer trials per condition.

Since VL-DBS data was not available for monkey N, the resulting latent space of the newly trained VAE was less structured compared to the one derived from the data of monkey T, which captured multiple functional trajectories. Instead, the latent space for monkey N appeared to encode changes predominantly along a single direction, defined by the anesthesia and the 5V CT-DBS clusters. Still, in the same fashion as with the other monkey, we found potential targets that, when perturbed, led the dynamics of the system to the regions occupied by the experimental conditions. In particular, as with monkey T, perturbation to the PFCm led the dynamics toward the 3V CT-DBS cluster, and perturbation to the PMCvl toward the 5V CT-DBS cluster (**Figure S6**). In contrast, HC perturbation displaced the dynamics to a region of the latent space uninhabited by any experimental conditions – whereas in monkey T it kept the dynamics inside the cluster of the anesthesia condition. **Figure S7** quantifies these perturbations by showing the distributions of distances from the 20% closest perturbation simulations to each experimental condition cluster.

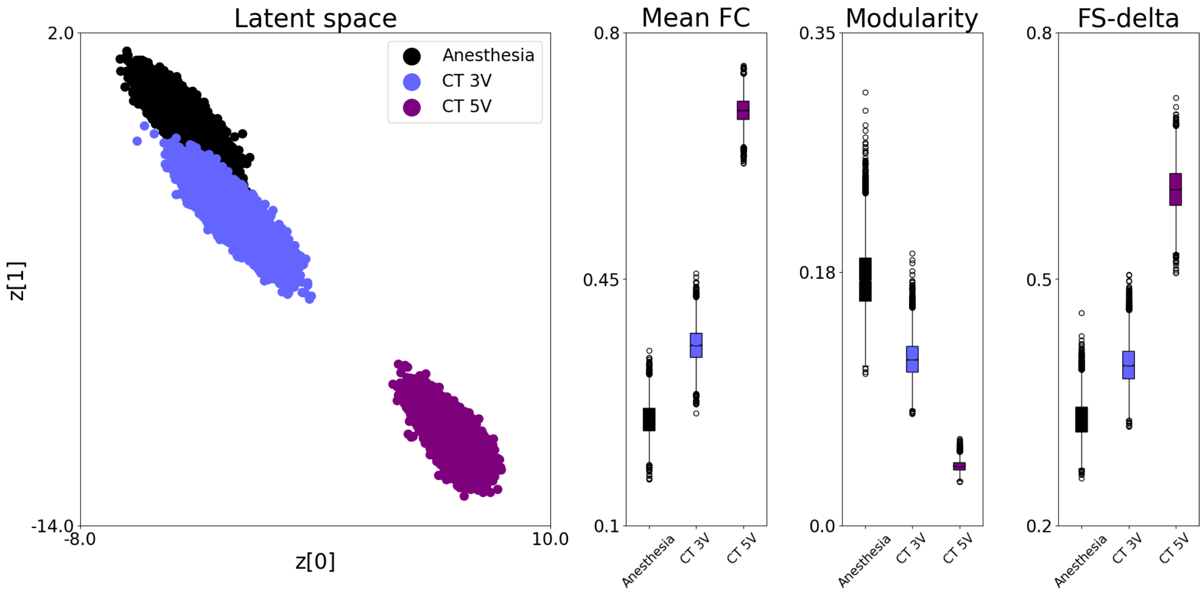


**Figure S4. Monkey N latent space organization and condition-specific metrics.** (Left) Depiction of the latent space with the three conditions encoded. (Right) Boxplots of the mean FC, modularity and FS-delta across conditions. All of the Kruskal-Wallis comparisons between distributions are highly significant ($p$ < *0.001*) after Bonferroni correction. Figure S5 contains Cohen's d metrics for said comparisons.

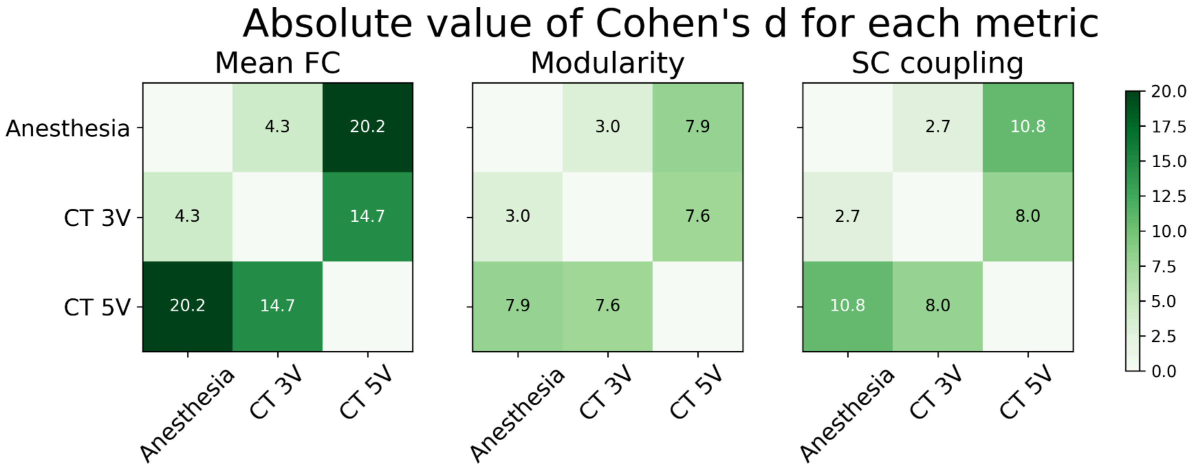


**Figure S5. Effect sizes for metric comparisons between the conditions for monkey N.** Cohen's d values for each metric, comparing distributions between conditions. This complements the results shown in the right panels of Figure S4.

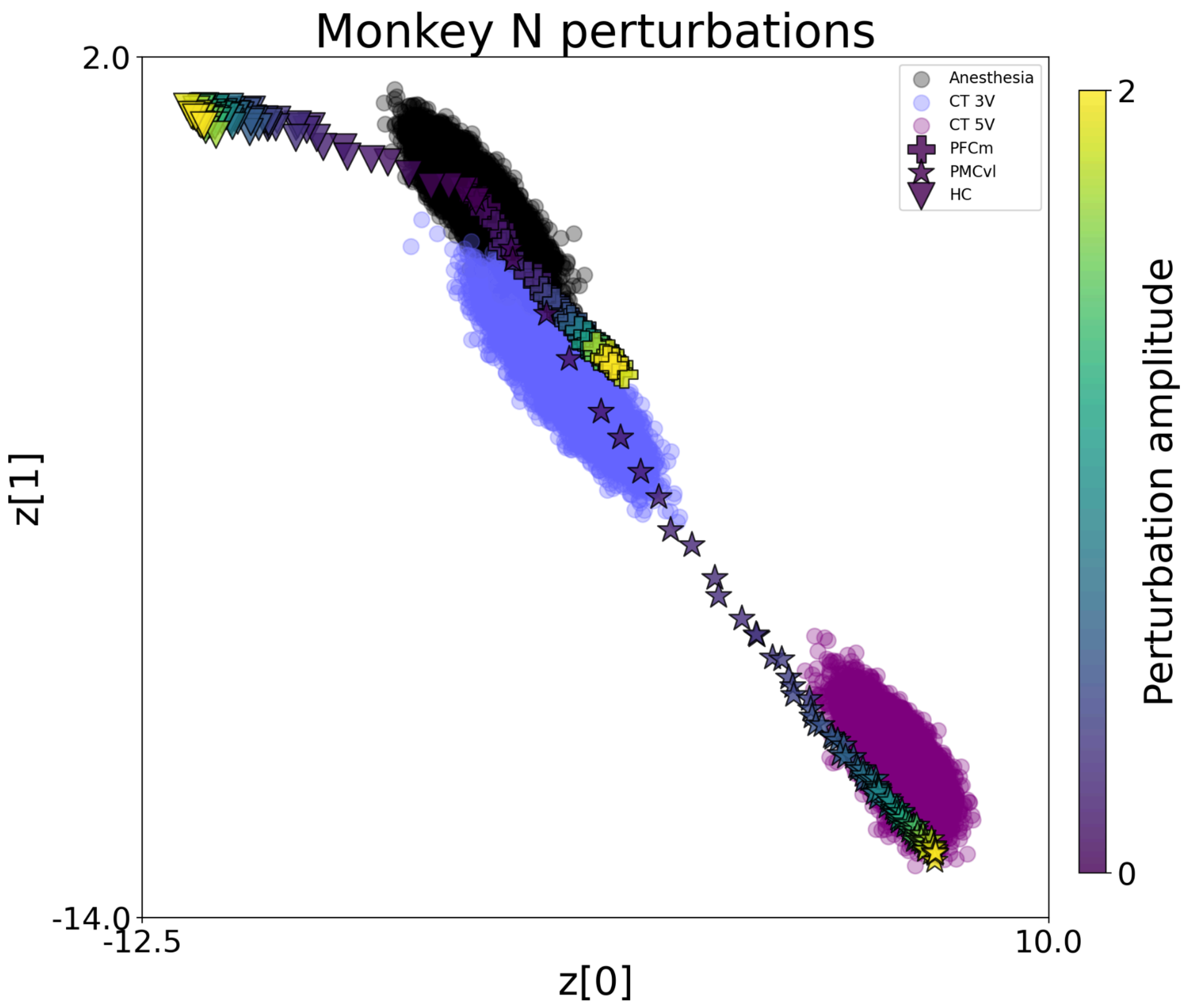


**Figure S6. Latent space trajectories for different perturbation targets in monkey N.** PFCm perturbation led the system toward the 3V CT-DBS dynamics and PMCvl perturbation shifted dynamics toward the 5V CT-DBS cluster, consistent with the results for monkey T. In contrast, HC perturbation moved the dynamics to an area of the latent space that is not related to any experimental condition –whereas in monkey T, HC perturbation did not generate a trajectory outside of the space covered by the anesthesia condition

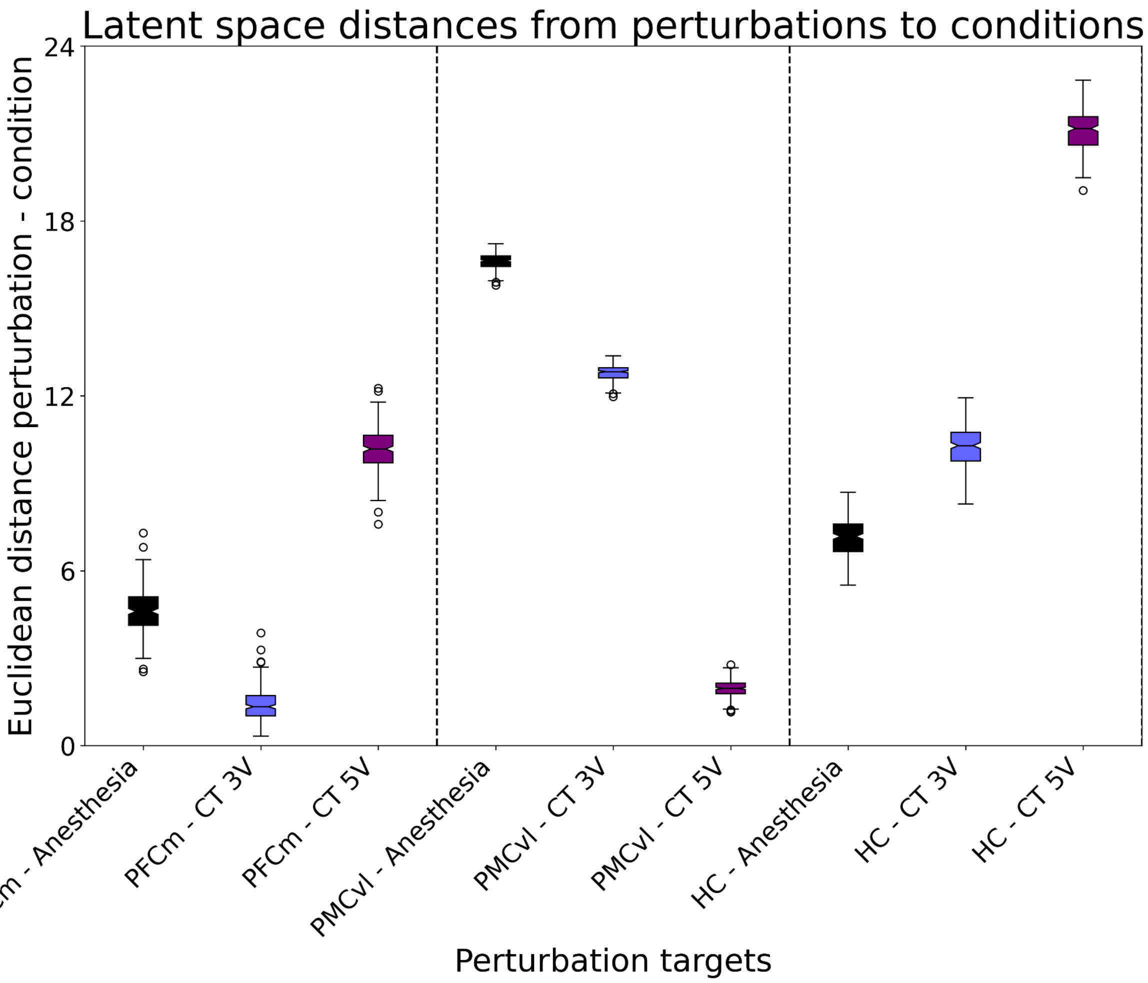


**Figure S7. Latent space distances between perturbed dynamics and the different conditions.** For each target, we chose the 20% of simulations that get closer to each of the experimental conditions' clusters. All comparisons were highly significant after correction ($p < 0.001$). This figure complements Figure S6, showing that perturbation targeting PFCm drove the dynamics closer to the 3V CT-DBS stimulation dynamics, PMCvl perturbation toward 5V CT-DBS stimulation dynamics, and HC perturbation away from all experimental conditions.
